# Force Induces a Collective Inactivity in Dyneins to Auto-Regulate Bidirectional Transport of Vesicles

**DOI:** 10.64898/2026.10.08.757518

**Authors:** Dwiteeya Chaudhury, Aniket Kumar, Paulomi Sanghavi, Sudipto Muhuri, Mithun Mitra, Roop Mallik

## Abstract

The back-and-forth (bidirectional) motion of Endosomes and Phagosomes along microtubules is fundamentally important for nutrient absorption, receptor recycling, pathogen degradation and many other downstream cellular pathways. Bidirectional motion requires opposing force from the kinesin and dynein motors, causing intermittent Tug-of-Wars (ToWs) followed by “runs” where winning motors drive their cargo until the next ToW. Dynein displays a counter-intuitive behaviour of catch bonding tightly to the microtubule against opposing force, but why such a response is needed remains unknown. Here we force dynein to catch-bond using an optical trap when dynein-teams are transporting phagosomes *in-vitro* and inside cells. An unexpected form of *<u>collective</u>* inactivity emerges, where larger dynein-teams remain inactive for longer after they have been catch-bonded. Kinesin exhibits a high probability of reversing the cargo during such periods of inactivity of dyneins. Therefore, dynein’s inactivity allows cells to auto-regulate the effect of opposing molecular forces for the purpose of achieving bidirectional motion of endosomes/phagosomes. We propose that an inactivating conformational change is induced when dynein detaches against strain from its catch-bonded state. Collective inactivity follows because each inactive dynein exerts an entropic force on the other to delay the entire team from being activated. We capture the dynamics of this process through a statistical model of Entropically Constrained Reactivation, obtaining predictions in excellent agreement with experimental data. The emergent behaviour observed here possibly prevents a “runaway dynein” situation inside cells, its simple entropic origin suggesting broader relevance to catch-bonded receptor ligand interactions operating under conditions of cellular crowding.

## INTRODUCTION

Cellular functions require diverse membrane-bound organelles to be delivered to specific locations inside cells. This delivery requires the back-and-forth (bidirectional) motion of organelles that are carried as “cargoes” along microtubules (MTs) by the kinesin and dynein motor proteins (1–3). Kinesin and Cytoplasmic dynein (hereafter dynein) generate force respectively towards the plus and minus direction along MTs, and may engage in Tug-of-Wars on cargoes as they move around in back-and-forth manner (4–6). Bidirectional motion is ubiquitous inside cells, being observed for many organelles (2, 3, 7–9) and can be reproduced *in-vitro* for lipid-bound vesicles (10) or DNA scaffolds (11) attached to dynein and kinesin. The mechanisms that generate bidirectional motion are of fundamental importance to cell biology (1–3, 12), inspiring a vast body of experimental and theoretical work which attempts to explain the opposing activity of dynein versus kinesin on a cargo through the Motor-Coordination or Tug-of-War (ToW) models. The coordination model posits that adaptor proteins bind/unbind to motors to regulate motor activity and cargo transport (1, 12, 13). In contrast, the ToW model posits that cargo-bound kinesins and dyneins engage in intermittent ToWs to drive motion depending on their force, numbers, geometry and attachment/detachment kinetics on the cargo (1, 6, 11, 14).

Early endosomes and phagosomes exhibit approximately equal rapid motion in plus and minus directions (called “runs”) that are interrupted by ToWs and reversals in direction (see Fig 1A), thus yielding little ***<u>net</u>*** motion of the cargo (4, 6, 15, 16). This process can be described by the tossing of a hypothetical coin, where force-generation by one class of motors is followed with equal probability of force-generation by the same or the opposite class (17). Adaptor proteins and/or membrane lipids (e.g. cholesterol) can bias this coin, thus modulating cargo localization over longer timescales(18). Recently, a helical domain in the mitochondrial adaptor protein TRAK was proposed as a molecular switch that coordinates kinesin versus dynein driven transport of mitochondria (12). Note, however, that the leading and lagging ends of mitochondria exhibit opposing motion (stretching) as it moves inside cells (5). These shape-changes in mitochondria are quite similar to endosomes, where ToWs are well established to drive bidirectional motion(6). We therefore believe that the Coordination and ToW models can be unified, where adaptor proteins (e.g. TRAK, dynactin) act as “coordinating molecules” that modulate motor ON/OFF rates and force-generation to change the outcome of ToWs, and therefore the nature of cargo motion by (18, 19). How individual motors and motor-teams respond to opposing force assumes primary importance in this picture of bidirectional transport.

**Figure 1.**
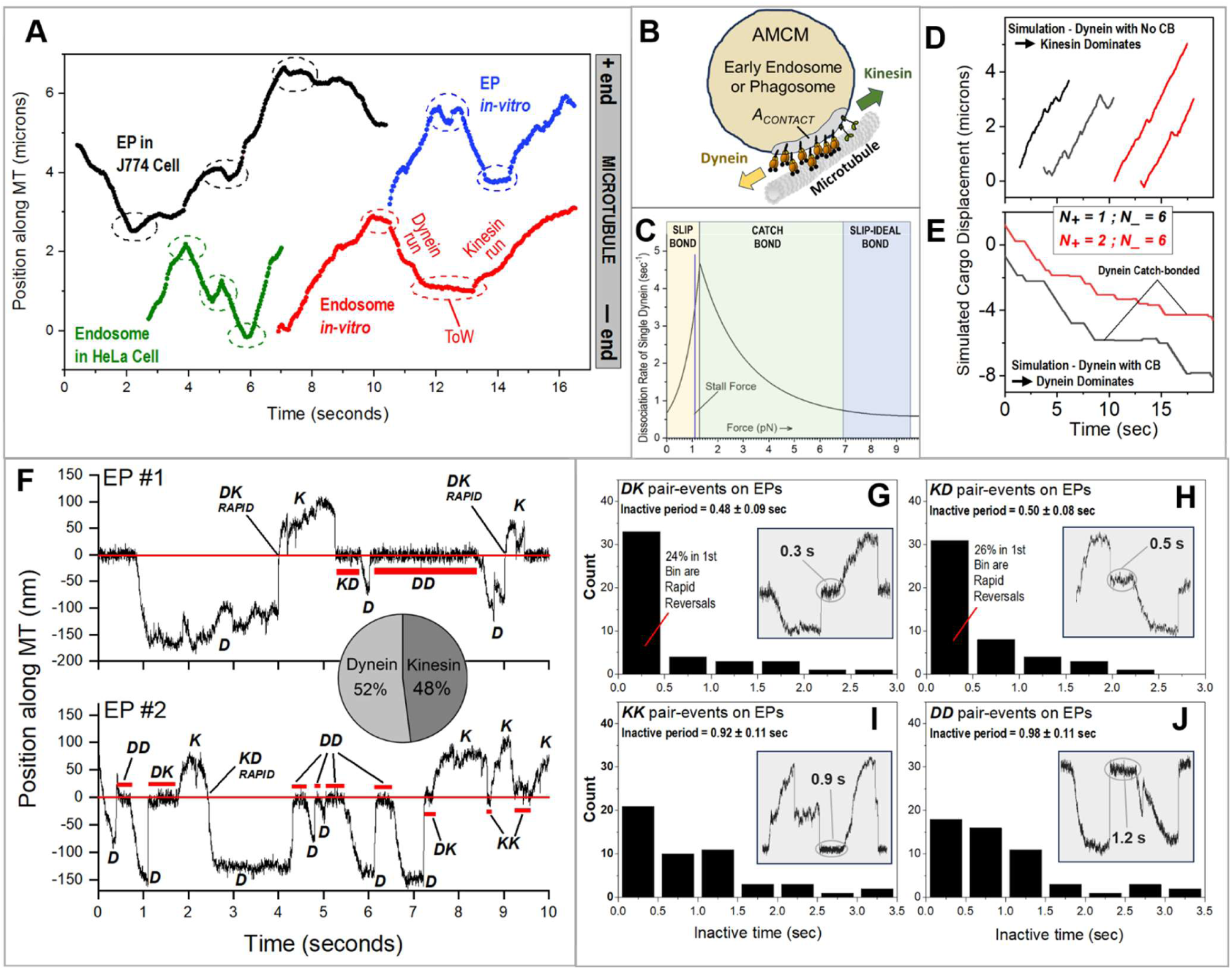
Activity and Inactivity of Opposing Motors during Bidirectional Motion. A) Representative video tracks of bidirectional (back-and-forth) motion of early phagosomes (EPs) and Endosomes along microtubules (MTs) inside cells and in an *in-vitro* assay. MT orientation is shown on the right. B) Schematic of an early phagosome (EP) depicting the Asymmetric Motor Competition Model (AMCM), where 6-7 dyneins (orange) and 1-2 kinesins (green) attach to the MT from an area of contact (*A_CONTACT_*) on the EP. The opposing force-vectors for both motors are shown (arrows), signifying the possibility of Tug-of-War (ToW) between motors. *C)* The Slip/Catch/Slip-Ideal bond behaviour of a single dynein motor as a function of increasing load force (see main text for more details). Figure based on *Kunwar et. al. PNAS* (*2011*) and *Puri et. al. Phys. Rev. Res.* (*2019*). D) Monte Carlo simulation of cargo motion for the indicated number of kinesin (*N*_+_) or dynein (*N*_–_) motors shown using position-time plots. Dyneins are modelled with no catch-bond (CB), resulting in motion dominated by plus-directed kinesins. E) As in (D), but dyneins now modelled allowing for a catch-bond according to Fig 1C, resulting in only minus-directed motion and paused states. Neither panel (D or E) reproduces the back-and-forth motion seen for endosomes/phagosomes seen in Fig 1A. F) Force generation events from Dynein (***D***) and Kinesin (***K***) against an optical trap on two representative EPs (EP#1 and EP#2). The trap centre is denoted by a red line. The inactive periods occurring within the four possible kinds of pair-events (***DK, KD, KK, DD***) have been indicated (thick red lines along X-axis). Some reversals are marked “RAPID” when the intervening inactive period is < 0.1 sec. These can involve ToWs between motors (see main text). **<u>Inset</u>**:– Pie chart illustrates overall fraction of dynein (***D***) and kinesin (***K***) events across multiple (= 50) EPs. G) Histogram of inactive periods observed between a ***D*** event that is succeeded by a ***K*** event (i.e. within a ***DK*** pair-event). The mean (± SEM) of inactive period is mentioned. Inset represents a typical ***DK*** pair-event with the inactive period indicated. H) Histogram of inactive periods within ***KD*** pair-events. Inset represents a typical ***KD*** transition. I) Histogram of inactive periods within ***KK*** pair-events. Inset represents a typical ***KK*** transition. J) Histogram of inactive periods within ***DD*** pair-events. Inset represents a typical ***DD*** transition.

While Early Phagosomes (EPs) exhibit equal kinesin and dynein activity, the molar concentration of dynein on EPs is significantly higher than kinesin-1 (4, 20, 21). Fitting force-histograms to a sum of multiple Gaussian profiles suggests that dynein generates ∼1.1 pN force on phagosomes inside cells (22), on lipid droplets in *Drosophila* embryos (23), on purified phagosomes (18) and on dynein-coated beads (24–26). The lowest unitary force generated on EPs in minus direction was found to be ∼1.2 pN after partial inactivation of dynein (17). Kinesin (*Dictyostelium* Unc104 or mammalian kinesin-1) generates a force between 5-6 pN on phagosomes (6, 18, 27). We therefore believe that the bidirectional motion of early endosomes involves multiple (approx. 4 to 8) weak and detachment-prone dyneins engaging in force-balanced ToWs against a few (1 or 2) strong kinesins. This scenario is schematized in Fig 1B, and has been described as the *Asymmetric Motor Competition Model or AMCM* (6). The use of multiple dyneins likely enables cargo motion to be tuned by controlling dynein (but not kinesin) number, as indeed observed on melanosomes (28) and endosomes (29).

Of particular relevance therefore is to understand how team-function emerges from the known single-molecule biophysical properties of motors. For example, can motors within a team cooperate (30, 31)? Evidence for such cooperation was seen when the magnitude and duration of force generated by an *N*-dynein team increased linearly with *N* (22). The mean first passage time before a dynein-team detaches against load, i.e. the dynein-team’s tenacity, was calculated by modelling the attachment/detachment of dynein as a Brownian particle (32). These authors concluded that its catch/slip-ideal bond behaviour, hereafter referred to as catch-bond, reduces dynein’s unbinding rate at high-load regimes (see Fig 1C). The special ability of a single dynein motor to catch-bond to the MT has been quantified in detail (33). This catch-bond may cause the force and tenacity of dynein-teams to increase linearly with their size, perhaps why cells assemble dynein-teams of different sizes tuned to specific cellular functions (31). It has also been suggested that dynein’s catch-bond can explain (34) why inhibiting one kind of motor (dynein or kinesin) blocks transport in both directions (1, 7).

Here we ask if dyneins within a team get catch bonded during ToWs against kinesin, and if so, how the catch-bond affects cargo motion ***<u>AFTER</u>*** a ToW. Our findings suggest that dynein gets inactivated when it detaches from the catch-bonded state induced by force from kinesin during a ToW. Importantly, when such catch-bonds are imposed on a team of dyneins, each inactive dynein hinders the other from getting activated. This collective response therefore “silences” a dynein-team for a certain duration after it has generated force, allowing kinesins to engage and achieve a balance of plus-versus-minus cargo motion. We rationalize this “collective inactivity” of dyneins through a simple mathematical model of *Entropically Constrained Reactivation*. These findings may have relevance other mechanobiological processes involving catch bonded receptor-ligand interactions – for example cell migration, wound healing, cell-cell and bacteria-substrate adhesion, sensing of acoustic signals in the ear etc.

## RESULTS

### TUG-OF-WAR MEDIATED BIDIRECTIONAL MOTION OF ENDOSOMES AND PHAGOSOMES

Fig 1A shows representative displacement versus time records of single Phagosomes and Endosomes moving along microtubules (MTs) inside cells and in reconstituted assays *in-vitro*. The intracellular early phagosomes (EPs) consist of a latex bead (750 nm diameter) phagocytosed inside a J774 mouse macrophage cell and visualized within 10 minutes after it was ingested [See Methods and Ref. (22)]. Displacements of the EP towards cell periphery or cell centre are respectively identified to be plus (kinesin-driven) or minus (dynein-driven) movements with high confidence. Supplementary Video S1 shows the motion of EPs inside a J774 cell. Latex beads can also be phagocytosed into *Dictyostelium* cells to prepare and isolate EPs which exhibit vigorous bidirectional motion when placed on *in-vitro* polymerized MTs (6). The recording of *in-vitro* motion of an EP has been reported by us in Ref. (18). MTs labelled at the minus-end allow us to identify the direction of *in-vitro* motion as plus or minus-directed (35). As seen in Fig 1A, the motion of purified EPs is qualitatively and quantitatively similar to cargoes moving inside cells [see Ref. (6) for a detailed analysis]. EPs and endosomes exhibit segments of fast plus and minus motion (“runs”) with almost equal probability and equal run-length, resulting in little **<u>NET</u>** motion of these cargoes. These runs are interrupted by low-velocity “paused” states (see Fig 1A) that may or may not be followed by a reversal. The paused states involve physical extension of endosomes along the MT, and are thus identified as ToW-states (6). Motors cannot deform EPs because they enclose rigid beads, but the similar duration and frequency of pauses makes a strong case for ToWs on EPs (17).

ToW-mediated bidirectional motion of cargoes has been investigated extensively using computer simulations. Muller *et. al.* discussed motion driven by the same number [*N+* = *N* = *N–*] of kinesin and dynein motors (14). Such a situation is very unlikely on EPs and endosomes, where a few “strong” kinesins [*N+* = 1 to 2; Stall force ∼6 pN] engage in force-balanced ToWs against many “weak” dyneins [*N–* = 6 or more; Stall force between 1 to 2 pN]. This situation was discussed earlier as the AMCM (6, 22, 31) that is schematized in Fig 1B. The time-evolution of such an “asymmetric” system of motors [*N+* = 1 to 2 and *N–* = 6] was analyzed by incorporating the catch-bonding of dynein through a threshold force bond deformation (TBFD) model (34, 36). Figs 1D and 1E show the results of Monte Carlo simulations under unequal load-sharing by motors (Methods). Switching OFF dynein’s catch-bond in these simulations causes overwhelmingly plus-directed runs because the dyneins detach rapidly in a cascade during ToW against kinesins (Fig 1D). Switching ON the catch-bond caused long minus-directed runs with frequent pauses because catch-bonded dyneins lock on to MTs whenever they sense opposing force from kinesin, thus almost completely blocking plus-directed motion (Fig 1E). These simulations are in good agreement with earlier work based on the TBFD model (34, 36). *How then do EPs, endosomes, mitochondria and other cellular cargoes maintain an equal balance of plus and minus directed runs interspersed with ToWs, resulting in little **<u>net</u>** motion of the cargo?* Clearly, reality lies somewhere in between. Dynein does require a catch-bond to resist kinesin, but the catch-bond must be modulated in some way so that kinesin also gets an equal chance to drive cargo motion.

### ANALYSING BIDIRECTIONAL MOTION OF PURIFIED EARLY PHAGOSOMES IN AN OPTICAL TRAP

Geometrical constraints mean that only motors residing within a certain area (= *A_CONTACT_*; see Fig 1B) on the spherical EP can attach to the MT to generate force (17, 18, 22). Fig 1F shows the motion of two purified EPs (EP#1 and EP#2) held above a MT using an optical trap, where the trap centre is at zero position (red line). Supplementary Video S2 shows the recorded motion of an EP against the optical trap. Optically trapped EPs allow observation of repeated motor-driven motion of a single cargo at high spatial and temporal resolution using a quadrant photo-detector (Methods). Switching off the trap causes these EPs to exhibit long plus and minus runs similar to Fig 1A. However, we prefer trap-based experiments to interrogate bidirectional motion because obstruction from other cargoes and restrictive MT lengths severely limit the observation period for EPs when they move freely (i.e. without a trap). Excursions of the EP against the trap towards MT plus or minus ends are depicted respectively as ***K*** (kinesin) or ***D*** (dynein) events (Fig 1F). Agreeing with the nature of bidirectional cargoes (Fig 1A), ***K*** and ***D*** events occur with almost equal probability in the optical trap when counted across many EPs (Inset, Fig 1F). The trap allows easy identification of the inactive period between two successive ***K*** or ***D*** events. At this time, no motor generates measurable force and EPs reside at the trap centre. All motors have likely detached during this inactive period because the standard deviation (S.D.) in position of trapped EPs in a direction orthogonal to the MT during these inactive periods is same as S.D. of an EP trapped away from a MT (25).

For the analysis to follow, two successive events (i.e. pair-events) are more relevant than individual (***K*** or ***D***) events. Four kinds of pair-events can occur (***KD, DK, KK*** and ***DD)*** with an inactive period possible within each. The ***KD*** and ***DK*** pair-events involve a reversal in between, but ***KK*** and ***DD*** do not (Fig 1F). Figs 1G-1J show the distribution of inactive periods for all pair-events on EPs. The inset within each panel shows a representative pair-event of the respective type, with its inactive period mentioned. The mean inactive period (± SEM) across multiple EPs is also mentioned for every panel (Figs 1G-1J). The inactive period is <0.1 sec for ∼25% of the reversals, with some such examples denoted as ***DK*** *(RAPID)* or ***KD*** *(RAPID)* in Fig 1F. These rapid reversals involve opposing motors engaged simultaneously or in rapid succession to generate ToWs, as discussed earlier (17). These *RAPID* reversals are counted within the first bin of the histograms.

### PAIR-EVENTS INVOLVING REVERSALS HAVE SMALL INACTIVE PERIODS

Bidirectional stalls (Fig 1F) may help understand how motors are harnessed for endosome/phagosome fission, receptor recycling etc. Generating bidirectional motion, however, is not trivial as we realized during experiments with spherical supported lipid bilayers (SSLBs). SSLBs can be prepared in the form of beads that are enclosed by an artificial lipid membrane (37). When incubated with an ATP-releasate from rat brain, the total number of SSLBs moving in plus direction closely equals the number moving in minus direction. However, unlike EPs, an individual SSLB rarely reverses direction when it is held in the trap or when it is moving freely. If the same SSLB is pulled away from the MT using the trap, released briefly and then trapped again to be placed back onto the MT, then it has a ∼50% chance of moving in the opposite direction (37). Considering their ON/OFF rates, a diffusing motor (dynein or kinesin) that enters *A_CONTACT_* has only ∼1% probability of MT-binding before it diffuses away from the MT, as discussed in detail earlier (17, 37). Rotation of the cargo and/or diffusion of motors within the membrane of the cargo therefore do not appear to have a major effect, and by-and-large the same set of motors generate force repeatedly during experiments such as those depicted in Fig 1F.

Contrasting with SSLBs that rarely reverse direction, the frequent reversals in direction of early endosomes and EPs suggest that individual ***K*** and ***D*** motors are somehow kept in close proximity on the endosome/EP membrane to promote ToWs and reversals. This proximity likely derives from adaptor proteins that interact with both motors to assemble kinesin-dynein (***K-D***) complexes on a cargo membrane (12, 38–40). ***K-D*** complexes are unlikely to be present on artificially prepared SSLBs, which are therefore unable to reverse direction in the trap. One can write the following rate-equation for motors on an EP (17):-

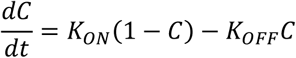

Here *K_ON_* and *K_OFF_* are the measured single-molecule attachment and detachment rates of ***K*** or ***D*** motors to the MT (Supplementary Table 1). Using the average numbers of motors residing within *A_CONTACT_* and the above rate equation, one finds that ∼6 dyneins would bind to the MT within ∼0.5 sec and 1 kinesin would bind within ∼0.3 sec to generate force (17). Thus, most inactive periods between ***K*** and ***D*** events should have a duration < 0.5 secs. Indeed, the inactive period between ***DK*** and ***KD*** pair-events follows exactly this behavior, with highest counts seen within the first bin (See Figs 1G and 1H).

A recent study measured the MT-rebinding rate of kinesin-1 while varying the bead-to-MT distance (*d*_B–MT_) in an optical trap (41). For *d*_B–MT_ ≈ 50 nm, a single kinesin-1 required ∼4 sec to rebind the MT. However, lowering *d*_B–MT_ to ≈20 nm caused fast rebinding within ∼0.3 sec. Similarly fast rebinding (within ∼0.2 sec) was seen in MT-gliding assays where multiple kinesins hold MTs close to the coverslip to make *d*_B–MT_ small (≈17 nm) (42). These observations strengthen the argument that cargo reversals involve ***K-D*** molecular complexes, because even when one class of motors detaches after a ToW, the other attached class would keep the cargo close to the MT to keep *d*_B–MT_ small. The unbound opposing motors can now rebind with faster rates to promote cargo reversals. This effect explains the smaller inactive periods between ***DK*** and ***KD*** pair-events (Figs 1G, 1H). Note that during free bidirectional motion of EPs (no optical trap; Fig 1A), an EP is held close to the MT (*d*_B–MT_ small) by one or the other class of MT-bound motors. We have therefore used the faster rebinding rate for kinesin when conducting the MC simulations. This rebinding rate is ∼5 sec^−1^ (Rebinding time = 1/5 ∼ 0.2 sec), see Supplementary Table 1.

### PAIR EVENTS WITH INVOLVING NO REVERSALS: DYNEIN REBINDS FASTER THAN KINESIN

The mean inactive period between ***KK*** pair-events that involve no reversals (Fig 1I) was significantly longer than the mean inactive period between reversals (***DK*** and ***KD*** pair-events; Figs 1G and 1H). Because kinesins are sparse on the EP membrane (Fig 1B), most ***KK*** pair-events likely involve two successive force generation attempts by the same kinesin. Indeed, the mean inactive period of 0.92 sec between ***KK*** pair-events on EPs (Fig 1I) is well within the range of single molecule re-binding times for kinesin discussed above (41). To verify whether this re-binding time is also applicable to our experiments, we purified DdUnc104, the dimeric kinesin that drives EP motion in *Dictyostelium* (6). DdUnc104 was coated onto beads and successive stalls were recorded at the single-molecule limit where <30% of beads moved (Supplementary Fig 1A). These experiments measured a mean rebinding time of ∼1 sec (Fig 2A), close to the mean inactive period of 0.92 sec between ***KK*** pair-events on EPs (Fig 1I). Optical trapping of purified mammalian kinesin-1 also yielded long rebinding times (mean = 1.3 sec, see Supplementary Fig 1B). All of these data suggest that the average rebinding time for a cargo-bound kinesin (DdUnc104 or Kinesin-1) is ∼1 sec in our optical trapping geometry.

**Figure 2.**
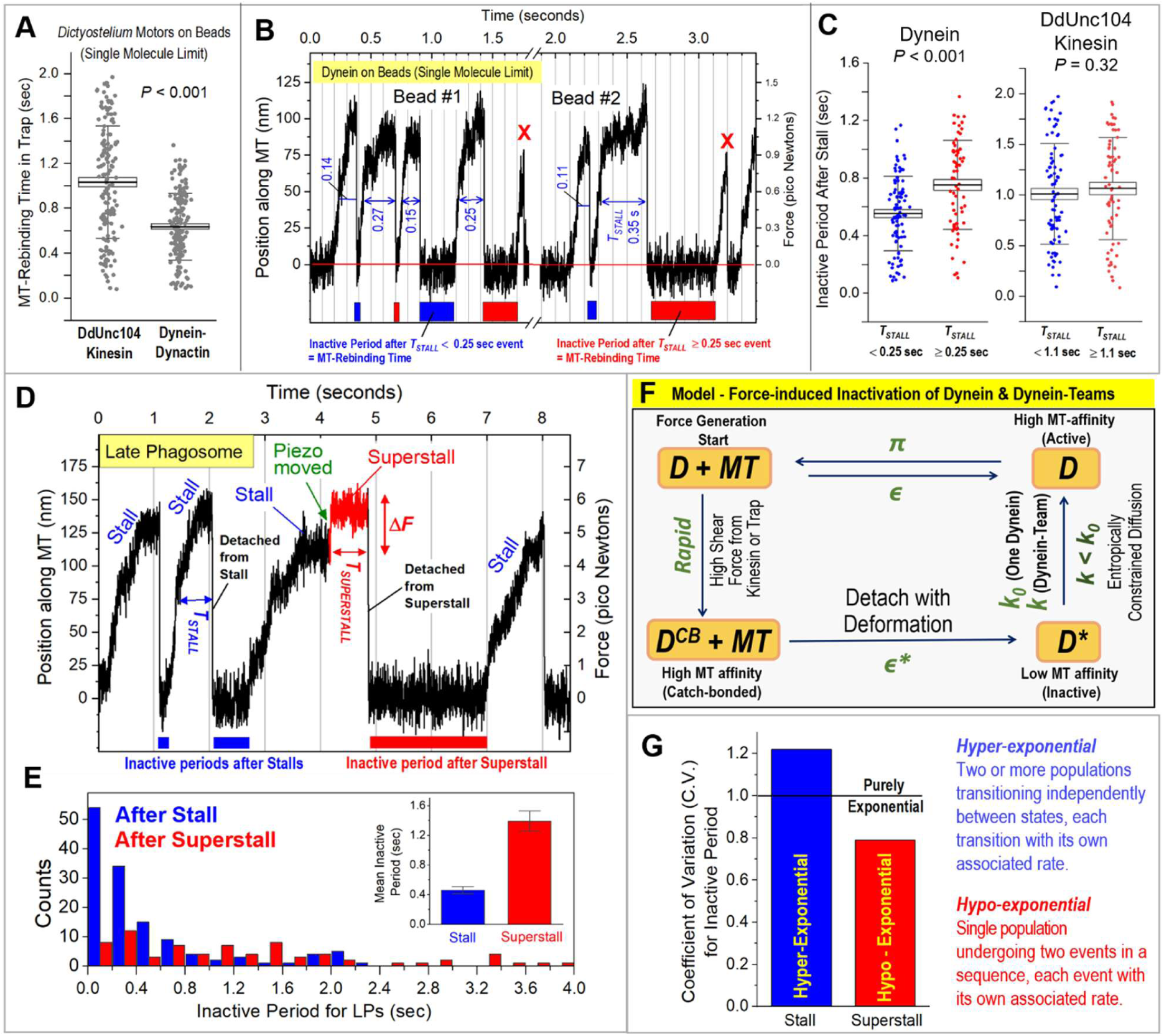
Dyneins Detaching from Catch-bonded State are Transiently Inactive. A) Time taken for *Dictyostelium* DdUnc104 kinesin and dynein-dynactin to rebind to a MT. Both motors were coated on beads. Optically trapped beads were held above a MT to assay rebinding. Experiments were done at single molecule limit (<30% probability of binding to MT in a 15-second period). Boxes are SEM and whiskers are SD. Mean is also indicated. B) Motion of two dynein-coated beads (Bead #1 and Bead #2) against an optical trap in the single-molecule limit. Force generated is also shown on the right axis (*F* = *Trap Stiffness* × *Position along MT*). Red line indicates trap centre (Zero position). Stall durations (*T_STALL_*) are indicated. Inactive periods after stalls of (*T_STALL_ <* 0.25 sec) or (*T_STALL_* ≥ 0.25 sec) are indicated respectively by blue and red rectangles. Red crosses are events that end in premature detachments (no plateau-like region). These events were not considered as stalls. Inactive period before a crossed event is thus considered for analysis, but the inactive period after a crossed event is not considered. C) The Inactive periods (i.e. time taken to rebind MTs) for dynein-dynactin and DdUnc104 kinesin, as measured in the experiments of Fig 2A are divided into two categories. For dynein:– ***(i)*** Inactive period after stalls of short duration (*T_STALL_* < 0.25 sec) and ***(ii)*** Of long duration (*T_STALL_* ≥ 0.25 sec). For DdUnc104 kinesin:– ***(i)*** Inactive period after stalls of short duration (*T_STALL_* < 1.1 sec) and ***(ii)*** Of long duration (*T_STALL_* ≥ 1.1 sec). See main text for the choice of this threshold. Boxes are SEM and whiskers are SD. Mean is also indicated. D) Force generation by a team of dyneins on a Late Phagosome (LP) purified from *Dictyostelium*. The measured stall force (right axis) suggests that a team of five to six dyneins is generating force. Piezo movement (indicated) induces an additional Superstall Force (Δ*F* = 0.3 × *F*_STALL_) on the dynein-team when it is stalled at a force *F* = *F*_STALL_ against the trap. The inactive periods are then measured between a stall event and the next event (blue; bottom axis), or between a superstall event and the next event (red; bottom axis). E) Histogram showing the distribution of inactive periods measured on LPs after Stalls (Blue; n = 138) or after Superstalls (Red; n = 76). Inset shows the mean inactive period for both cases (bars are SEM). F) Proposed Inactivation Model for Single Dynein or Dynein-Team. A single Dynein is shown in four states (***D***) Active, but not bound to MT. (***D+MT***) Bound to the MT and capable of taking steps/generating force (active). (***D^CB^+MT***) Catch-bonded tightly to the MT due to high shear force. (***D\****) Detached from the MT after catch-bond with a deformation that induces an inactive low-affinity state of dynein to MTs. The transition rates between various states are mentioned. A single dynein in ***D\**** state can get reactivated by returning to ***D*** state only by thermal diffusion (rate = *k_0_* for single dynein). For a dynein-team, this rate is further slowed down (*k < k_0_*) because of entropic constraints to diffusion. The rate *k* is described in terms of *k_o_* by the entropic constraint model (Equation 3). G) The coefficient of variation (*CV*) of the inactive period that was measured after Stalls or Superstalls. The line at *CV* = 1 corresponds to a purely exponential process. The *CV* for Stalls is > 1 but <1 for Superstalls, respectively signifying “Hyper” and “Hypo” exponential processes

In order to quantitatively compare the kinetics of kinesin versus dynein rebinding to MTs, we next coated beads with dynein-dynactin complexes purified from *Dictyostelium* (6), and recorded repeated force generation events at the single-molecule limit similar to the experiments with kinesin. The mean rebinding time of single dynein-dynactin to MTs was significantly smaller (∼0.6 sec) as compared to ∼1 sec for DdUnc104 kinesin measured under identical conditions (Fig 2A). The faster MT-rebinding of single dynein-dynactin complexes may derive from dynein’s floppy nature, large size and/or because dynactin provides an additional tether for this complex to bind the MT(43). This faster rebinding for single dynein, however, also poses a conundrum that motivates the present study. If multiple dyneins reside within *A_CONTACT_* on the EP (Fig 1B), then any one of them should be able to bind to the MT and rebinding should be significantly faster than a single dynein (= 0.6 sec). Unlike kinesin, multiple dyneins cooperate well to generate collective force as a team (22). Even if a few dyneins generate force, such an event would be detected by the quadrant photodetector used in our experiments [see Ref (17, 27, 37)]. *Why then do we see long inactive periods within **DD** pair-events* (Fig 1J)*?* This inactivity appears to be caused by the first force-generating event of dynein because ***KD*** pair-events, where kinesin generates force first, have short inactive periods.

### DYNEIN IS INACTIVE AFTER IT HAS RESISTED FORCE FOR A LONG DURATION

Fig 2B shows repeated stalls of a single dynein coated on a bead (in the form of a dynein-dynactin complex) that was purified from *Dictyostelium*. The tenacity of motors against opposing force can be measured from the *T_STALL_* (time spent above half-maximal force; see Fig 2B) in such experiments (22). A longer *T_STALL_* suggests that detachment was delayed (i.e. detachment rate was lowered), indicative of catch-bond formation against force from the trap. We will therefore use *T_STALL_* as a measure of the probability that a motor was catch-bonded during the stall. Because the mean *T_STALL_* = 0.25 sec for a single dynein (22), we divided the recorded stalls into two categories, namely [*T_STALL_* < 0.25 sec; *^T^*_STALL–SH0RT_] and [*T_STALL_* ≥0.25 sec; *^T^*_STALL–L0NG_]. We then measured the “inactive period”, i.e. the time elapsed between each *^T^*_STALL–SH0RT_ event and the next event (blue rectangles along X-axis in Fig 2B), or the time elapsed between each *^T^*_STALL–L0NG_ event and the next event (red rectangles along X-axis in Fig 2B). This inactive period was taken as the interval between two force-generation events where the cargo was within (3 × *SD*) distance from trap centre, *SD* being standard deviation in position of a bead trapped away from a MT (no force generation by motors).

These inactive periods (i.e. MT-rebinding times) are shown in Fig 2C for dynein, and are essentially the data for dynein shown in Fig 2A after division into the aforesaid *T_STALL_* categories. A significantly higher inactive period was seen for dynein after it had resisted the optical trap for *^T^*_STALL–L0NG_, likely getting catch-bonded in the process. Note that the force generated by dynein after a *^T^*_STALL–SH0RT_ event was the same as after a *^T^*_STALL–L0NG_ event (Supplementary Fig 1C). Dynein was therefore fully active once its inactive period had elapsed, regardless of whether it was previously catch-bonded or not. The *ON*-rate of dynein for MT-binding (*K_ON_* = 2.5 sec^−1^) suggests that a single dynein should bind to MTs within (1/*K_ON_*) = 0.4 seconds (Supplementary Table 1). The measured mean inactive period of 0.5 sec for the *^T^*_STALL–SH0RT_ category is in excellent agreement with this value of *K_ON_*. However, for the *^T^*_STALL–L0NG_ category of dyneins which were presumably catch-bonded, the inactive periods persisted for ∼0.8 sec on an average (Fig 2C). Thus, interestingly, a dynein exiting from catch-bond was inactive for an additional period of ∼0.4 seconds when compared to a dynein that was presumably not catch-bonded.

We next conducted the same analysis on multiple-dynein driven beads where most stalls measured between 3 to 6 pN ^(Supplementary Fig 2A)^, with a mean *^T^_STALL_* of 0.75 sec that was higher than the mean *^T^_STALL_* of single dynein. This is consistent with our finding that *T_STALL_* increases linearly with the number of dyneins (22). For reasons explained earlier, we again divided the multiple-dynein stalls into ^[*T*^*_STALL_* ^< 0.75 sec; *T*^_STALL–SH0RT_^]^ and [*T_STALL_* ≥ 0.75 sec; *^T^*_STALL–L0NG_] categories. As expected from our earlier discussion based on the rate-constant equation, the *^T^*_STALL–SH0RT_ category exhibited a mean inactive period of ∼0.5 sec, but for *^T^*_STALL–L0NG_ category a significantly longer mean inactive period was measured (1.1 sec; Supplementary Fig 2B). Thus, similar to single dynein, catch-bonding also appeared to induce a longer inactivity in dynein-teams. Once this inactive period had elapsed, dyneins appeared to be fully active because they generated the same force whether they had been catch-bonded (previous *T_STALL_* ≥ 0.75 sec) or not (previous *T_STALL_*<0.75 sec), see Supplementary Fig 2C.

Having observed that catch-bond formation (as inferred from longer *T_STALL_*) induces an inactive period in dynein, we tested for this behaviour in the DdUnc104 kinesin that engages in ToWs with dynein on endosomes/phagosomes (6). Similar to the experiments with dynein, repeated force-generation by single DdUnc104 coated on beads was recorded (Supplementary Fig 1A) to obtain a mean *T_STALL_* of 1.1±0.55 sec. The inactive periods between stalls were measured and divided into periods preceded by a stall of *T_STALL_* < 1.1 sec, or periods preceded by a stall of *T_STALL_* ≥ 1.1 sec. There was no significant difference in the inactive period between these categories (Fig 2C), a finding that contrasts with dynein. Thus, kinesin which is not known to catch-bond to the MT in optical trap assays, does not enter into a force-induced inactive state. Note again that just like dynein, kinesin could also generate force normally once the inactive period had elapsed, regardless of whether the previous event had short or long *T_STALL_* (Supplementary Fig 1D).

### SUPERSTALLS CONFIRM THAT CATCH-BOND FORMATION INACTIVATES DYNEIN TRANSIENTLY

Catch-bond formation is a single-molecule phenomenon that requires a molecular-level shear force which is difficult to replicate in bulk assays. The underlying structural changes that cause catch-bonding have been explored for only a few systems (44), and are not known for the dynein-MT bond (see Discussion). It was therefore not possible for us to generate dynein mutants defective in catch-bond formation. Even if this was possible, such a mutant could altogether prevent catch-bond formation and therefore not allow the investigation of a post catch-bond effect. Therefore, we took a biophysical approach to force catch-bonds within a dynein-team, and then measured the inactive period following catch-bond formation. We chose Late Phagosomes (LPs) purified from *Dictyostelium*which exhibit overwhelmingly minus-directed motion driven by multiple dyneins in an *in-vitro* motility assay (18). Fig 2D shows representative stalls of an LP against the trap. We indicate a time point during a stall where a piezo stage was moved in the direction of LP motion, thus exerting an additional superstall force Δ*F* = (0.3 × *F*_STALL_) on dyneins over and above the stall force (*F*_STALL_) generated by the dynein-team. Because the piezo stage was not moved during every stall, a typical experiment contained both *Stall* and *Superstall* events (see Fig 2D).

The *T_STALL_* under superstall conditions (i.e. the *T_SUPERSTALL_*) was significantly higher (Supplementary Fig 3A). This confirms that dynein-teams survived longer against the trap when superstalled, and therefore likely contained a larger fraction of catch-bonded dyneins. We next measured the inactive period after each stall or superstall to observe that superstalls clearly shifted the distribution towards longer inactive periods (Fig 2E), as also seen from the mean inactive period (Inset, Fig 2E). Thus, forcing catch-bonds within the dynein-team on LPs delayed the dyneins from generating force again. Applying a “high superstall” force [Δ*F* = (0.5 × *F*_STALL_)] did not further increase *T_SUPERSTALL_* (compare superstall versus high superstall data in Supplementary Fig 3A). Therefore, almost all dyneins had already been catch-bonded by the superstall force. Also note that the mean *F*_STALL_ was same for events following a stall or superstall (Supplementary Fig 3B). Dyneins therefore resumed their normal force-generation after an inactive period following superstall/catch-bond formation. Note that superstall experiments using DdUnc104 kinesin coated beads at single molecule limit showed that *T_SUPERSTALL_* << *T_STALL_* (Supplementary Fig 3C). This verifies the slip-bond behaviour of kinesin against force, as against the catch-bond response of dynein (Supplementary Fig 3A).

### STALL VERSUS SUPERSTALL: DISTRIBUTION OF INACTIVE PERIODS

The above biophysical data confirms that dynein-teams remain inactivated for a significant period after the dyneins detach from a catch-bonded state. We propose a ***“Dynein Inactivation Model”*** to explain this behaviour in Fig 2F. In this model, single dynein undergoes an unknown structural change when it detaches under strain from a catch-bonded state (i.e. during the ***D^CB^*** + ***MT*** → ***D***^∗^ transition). This force-induced deformation persists after detachment from the MT, trapping dynein in an ***off-pathway inactive state*** (***D* state***). The structural basis for this proposed deformation is elaborated further in the Discussion section. Dynein must now recover to its active state (***D***^∗^ → ***D***; see Fig 2F) via thermally-induced diffusion before it can again bind MTs to generate force. Recall that for single-dynein the (*T_STALL_*<0.25 sec) and (*T_STALL_*≥ 0.25 sec) categories respectively had mean inactive periods of ∼0.4 sec and ∼0.8 sec (Fig 2C). Because dynein was not catch-bonded in the former category, the ^(***D*** → ***D*** + ***MT***)^ transition required ∼0.4 sec. This value is in excellent agreement with the reported ON-rate (*π* = 2.5 *sec*^−1^; 1⁄*π* = 0.4 sec) of single dynein (Supplementary Table 1). For the latter category (*T_STALL_* ≥ 0.25sec), we propose that such dyneins must undergo two sequential events 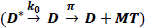 before they can again generate force. Because these dyneins were inactive for ∼0.8 sec, the ***D***^∗^ → ***D*** transition must also require ∼0.4 sec, yielding a rate *k*_0_ = 1⁄0.4 = 2.5 *sec*^−1^ for the ***D***^∗^ → ***D*** transition of a single dynein.

To move beyond single dynein and understand how the ***D\**** state impacts force generation by dynein-teams, we next analyzed the distribution of inactive periods after normal stalls and superstalls across a range of forces generated by dyneins on LPs. We assume that there is a higher fraction of ***D\**** dyneins in a dynein-team after superstall because excess force should induce catch-bonding with higher probability. After a dynein-team has detached, dyneins would attempt to bind the MT again, with each binding event governed by rate-limited exponential kinetics. When one or more dyneins bind and move (***D + MT*** state in Fig 2F), the inactive period ends, and is registered as cargo motion away from the trap centre. Thus, the distribution of inactive periods should contain information about the ***D\**** and ***D*** states, their relative populations within a dynein-team and the kinetics of conversion between these states.

The coefficient of variation for any variable is defined as 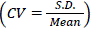, with *CV* = 1 for a purely exponential process. If *CV* < 1 then the underlying process that causes variation follows a *hypo-exponential* distribution where asinglepopulation undergoes two events in a sequence, each event having its own rate. However, if *CV* > 1 then the underlying process follows a *hyper-exponential* distribution characterized by two or more populations transitioning independently between states, each with its own associated rate. The above *CV*-based characterization of time intervals describes information transfer in telecommunication systems (45). We calculate that the distribution of inactive periods occurring after Superstalls for LPs has a *CV* < 1, but the inactive periods occurring after normal Stalls for LPs has a *CV* > 1 (Fig 2G). This difference suggests that Superstall causes almost all of the force-generating dyneins to catch-bond, thus creating a single homogeneous starting population of ***D\**** dyneins which then undergo two sequential transitions (***D\**** ◊ ***D*** and ***D*** ◊ ***D + MT***) to yield a hypo-exponential distribution of inactive periods. In contrast, normal stalls induce catch-bonds only in a fraction of dyneins, creating two different starting populations (***D*** and ***D\****) that transition independently to the (***D + MT***) state to yield a hyper-exponential distribution of inactive periods.

### FORCED CATCH-BONDING OF DYNEIN ALLOWS KINESIN TO REVERSE EARLY PHAGOSOMES *IN-VITRO*

We now return to our original question:– *Is bidirectional motion made possible inside cells because catch bonded dynein-teams remain inactive for a longer time, allowing kinesin to reverse the cargo?* To test this hypothesis, we first studied the bidirectional stalls of purified EPs on polarity labelled MTs *in-vitro*. Similar to experiments with LPs, we moved the piezo stage towards the MT minus-end during a dynein-driven stall of the EP to induce a Superstall (marked in Fig 3A), thus increasing catch-bond formation in dyneins. In between these superstall events, the piezo was not moved for many dynein-driven events to just record Stalls. We then scored whether the event following a Stall or a Superstall was again dynein-driven (no reversal) or kinesin-driven (reversal). Around 40% EPs reversed direction after Stalls, however ∼70% of the EPs reversed direction after Superstalls (Fig 3B). To test if this was a dynein-specific effect, a superstall force was exerted on DdUnc104 kinesin by moving the piezo towards the plus-end of the MT during the plus-directed stalls of EPs. Superstalling kinesin caused no increase in reversals, with ≈50% EPs reversing after stalls as well as superstalls (Fig 3B). Force-induced reversals therefore occur only when dynein is subjected to force, likely arising from its catch-bonding. This is expected because the kinesin-MT interaction shows slip-bond behaviour in contrast to dynein’s catch-bond (Supplementary Fig 3C).

**Figure 3.**
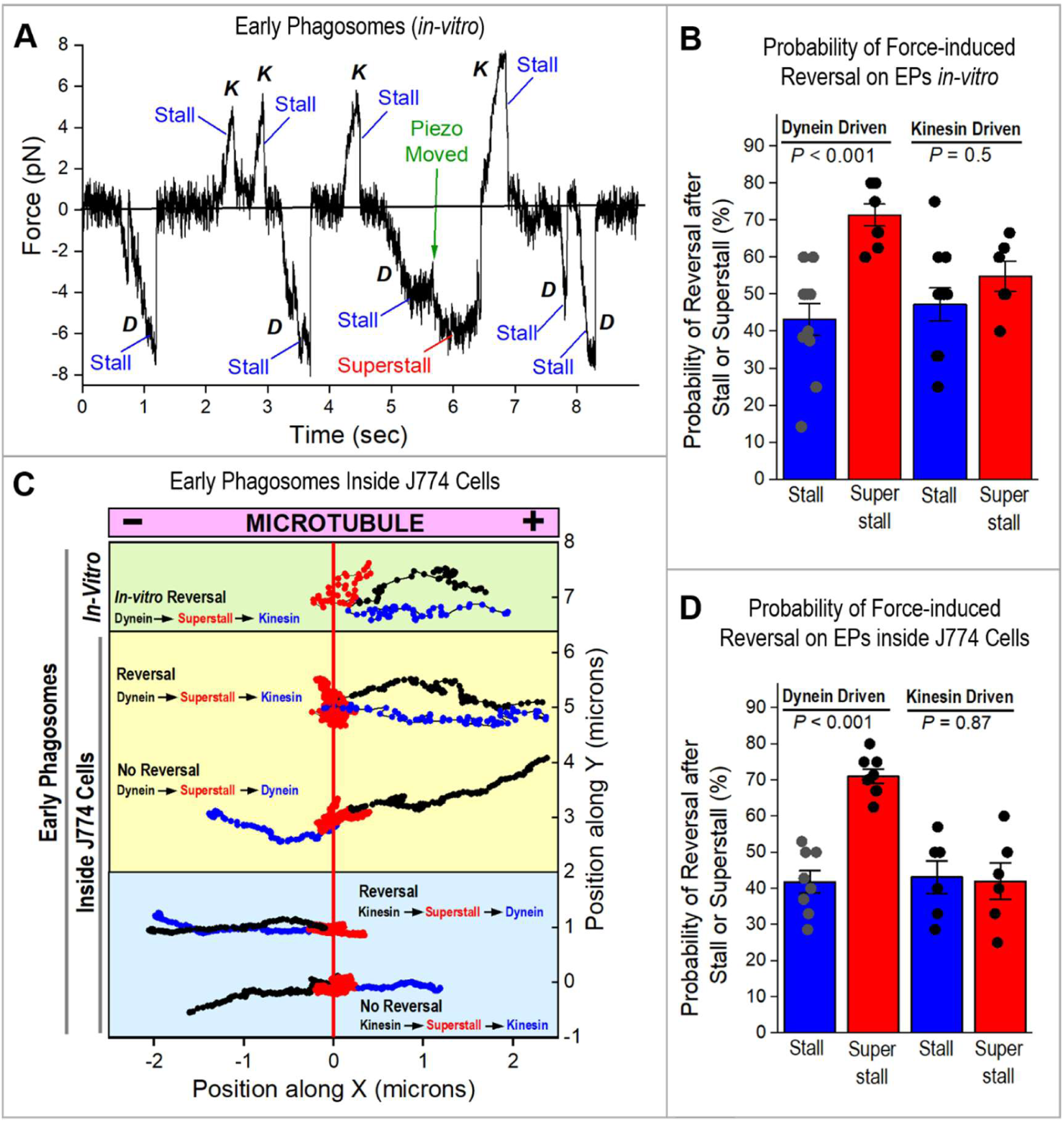
Kinesin has higher chance of Reversing the Cargo when Dyneins detach from Catch-bonded State. A) Bidirectional force-generation by Kinesin (***K***) or Dynein (***D***) on an Early Phagosome purified from *Dictyostelium*. Piezo movement (indicated) induces additional force to causes a Superstall on a team of ∼5 stalled dyneins. Note that there is a kinesin-driven event (reversal) right after the Superstall. B) Probability of reversal after a Stall (blue) or after a Superstall (red), as observed for EPs moving against an optical trap *in-vitro*. Data is shown where Superstall force was applied during dynein-driven motion, or during kinesin-driven motion. Superstalling dyneins increases cargo reversals, but such an effect is not seen after superstalling kinesins. Dynein-driven motion:– 55 EPs were tested for stalls and 30 EPs were tested for superstalls. Kinesin-driven motion:– 55 EPs were tested for stalls and 70 EPs were tested for superstalls. Error bars are SEM. C) *X*-*Y* position trajectories of Early Phagosomes (EPs) Superstalled *in-vitro* (green box) or inside J774 cells (yellow and blue boxes). Microtubule orientation is shown at the top. EPs were allowed to move freely, trapped at high laser power to induce superstall (red), then released to observe subsequent free motion that could occur in same direction as before trapping (no reversal), or in opposite direction (reversal). The superstall force was applied on dynein (green and yellow boxes), or on kinesin (blue box). See main text for more details. D) Probability of reversal after a Stall (blue) or after a Superstall (red), as observed for EPs moving against an optical trap inside J774 cells. Data is shown for events where Superstall force was applied during dynein-driven motion, or during kinesin-driven motion. Superstalling dyneins increases cargo reversals, but such an effect is not seen after superstalling kinesins. Dynein-driven motion:– Stalls (53 EPs tested across 7 biological replicates). Superstalls (47 EPs tested across 8 biological replicates). Kinesin-driven motion:– Stalls (35 EPs tested across 6 biological replicates). Superstalls (35 EPs tested across 6 biological replicates). Error bars are SEM.

### KINESIN REVERSES CARGOS MORE FREQUENTLY AFTER DYNEINS ARE SUPERSTALLED INSIDE CELLS

We next extended the superstall experiments inside J774 mouse macrophage cells that extend spindle-like projections with close-to-linear MTs within these spindles (Supplementary Fig 4A). We have used this system extensively to measure force from dynein-teams on late phagosomes that move predominantly in minus-direction (22). We pulsed cells with 750 nm (diameter) latex beads and observed the MT-dependent motion of so-formed phagosomes inside cells. To interrogate EPs, we shortened the chase time of phagosomes to 5 min after pulsing them for 5 min (Pulse + Chase = 10 min). We observed vigorous back-and-forth motion of EPs for upto ∼10 minutes after the chase inside cells (Supplementary Video S1). Unlike *in-vitro* assays, individual MTs are not visible and the precise MT orientation is unknown during a trapping experiment inside cells. Instead of moving the piezo to cause a superstall, we therefore rapidly increased the laser power to exert a higher force on EPs inside cells (see Methods). EPs were used to record stalls (no change in laser power), or superstalls where EPs stalled at ∼6pN before an additional force of ∼3pN was exerted. The recorded superstall of a dynein-driven EP and its subsequent reversal in direction of motion is shown in Supplementary Video S3.

Movement was video-tracked to obtain individual trajectories of EPs (Fig 3C; see the yellow and blue boxes). Fig 3C also shows the track of a purified EP that stalled against the trap and then reversed direction along a single MT *in-vitro* (green box). It is clear from the similar *Y*-displacements in green, yellow and blue boxes that EPs inside cells were also likely reversing direction along a single MT. The yellow box in Fig 3C shows data for two EPs that are initially driven by dynein (black dots). A superstall force was applied, (red dots), after which the trap was turned off and the subsequent motion of EPs was recorded (blue dots). The first EP in yellow box reversed direction after superstall, but the second EP continued to move towards the MT minus end (no reversal). Similar experiments were done for dynein-driven EPs subjected to normal stalls (with no increase in laser power, trajectories not shown). Fig 3D shows that the reversal probability of EPs for normal stalls is ∼40%, agreeing with the bidirectional nature of EP motion. However, superstalling dynein increased the probability of reversals by almost two-fold inside cells (Fig 3D) in a manner similar to *in-vitro* superstall data (Fig 3B).

We next tested whether superstalling kinesin enhances dynein engagement to increase EP reversals. Fig 3C (blue box) shows trajectories of two EPs that are initially driven by kinesin (black dots). The EPs are trapped and superstall force is applied on kinesin (red dots). The first EP reversed direction (blue dots) after superstall, but the second EP continued to move towards the cell periphery (no reversal). Superstall force on kinesin caused no change in the fraction of reversals across multiple EPs, which continued to be ∼50% of the total events (Fig 3D). The recorded superstall of a kinesin-driven EP and its subsequent reversal in direction of motion is shown in Supplementary Video S4.

Overall, it therefore appears that a fraction of dyneins gets catch-bonded and transiently inactivated during normal stalls against the trap. This fraction is such that it gives both dyneins and kinesins an equal chance to engage after dyneins have detached from the MT and the stall has ended (Figs 3B, 3D). The same situation should also occur during ToWs because the force-balance between dyneins and the trap is now replaced by a force-balance between dyneins and kinesins. Superstalling dynein using the trap tilts the balance in favour of kinesin (∼70% reversals; Figs 3B, 3D) and serves as an experimental tool to verify our hypothesis. However, this condition is likely not encountered during ToWs inside cells which are force-balanced to induce ∼50% reversals. This balance achieves approximately equal motion in plus and minus direction for early endosomes/phagosomes, so that they can engage in kiss-and-run interactions with other organelles over a longer duration to undergo programmed maturation (16, 46).

### ENTROPICALLY CONSTRAINED DIFFUSION OF CATCH-BONDED DYNEINS: THEORETICAL MODEL

The superstall experiments apply a force (*F*_STALL_ + Δ*F*) on dyneins, where Δ*F* = (0.3 × *F*_STALL_). Increasing Δ*F* to (0.5 × *F*_STALL_) did not further increase *T_SUPERSTALL_* (Supplementary Fig 3A). Thus, superstall induces a catch-bonded state (***D^CB^*** + ***MT***) and a subsequent inactive (***D\****) state in all/most of the dyneins for experiments on *N* = 1 to 6 dyneins. Fig 4A shows the mean inactive period after superstall on LPs, where the number of force-generating dyneins varies from *N* = 1 to ∼6. The value of *N* is inferred from the measured *F*_STALL_ assuming 1.1 pN force per dynein, and an additive nature of these forces (22). The inactive period after superstall is proposed to involve a population of ***D\**** dyneins that are undergoing (***D***^∗^ → ***D*** → ***D*** + ***MT***) transitions. As a simplest model we first assume that these two transitions happen independently for every dynein, and therefore any one of the *N* dyneins can transition with equal probability. Let us denote *k*_0_ and *π* as the experimentally measured single-dynein rates for these transitions (Fig 2F and Supplementary Table 1). In this “No Interaction Model”, the variation of the mean inactive period *T*(*N*) with *N* can be solved exactly (see Appendix-1 in Supplementary Information). This solution admits an upper bound as below:-

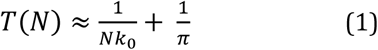

**Figure 4.**
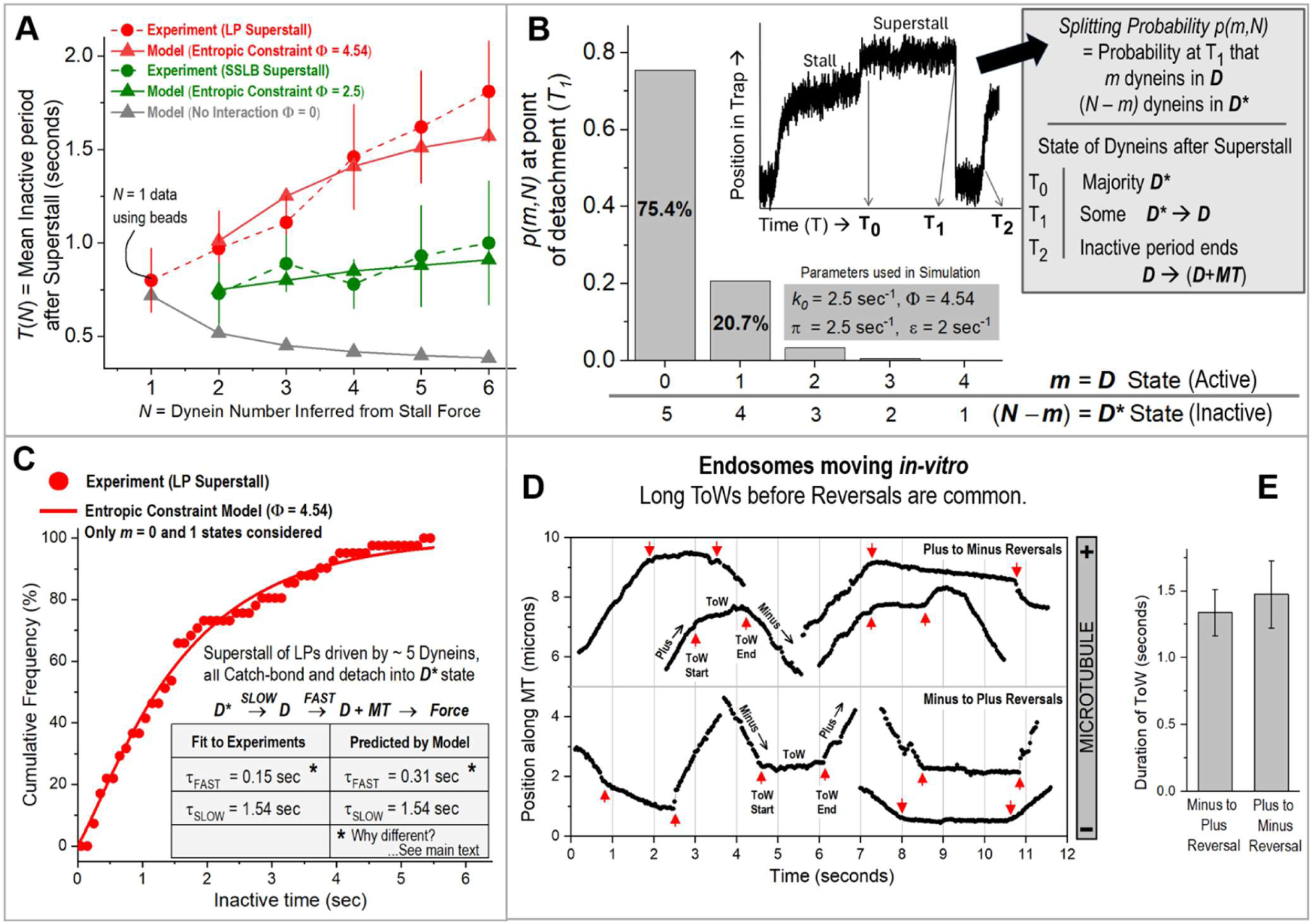
Reactivation Delay of Dynein-Teams after Catch-bonding and its Effect on Cargo Motion. A) The experimentally measured mean inactive periods *T(N)* for late phagosomes (LPs) and spherical supported bilayers (SSLBs) as a function of dynein number (*N*). *N* is inferred from measured stall force, assuming that 1.1 pN force corresponds to one dynein. The experimental data was obtained from superstall experiments done on beads (*N* = 1 dynein) or on LPs (*N* ≥ 2 dyneins). A single catch-bonded dynein must undergo 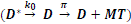 transitions before it can generate force again. These rates are taken as *k*_0_ = 2.5 sec^−1^ and *π* = 2.5 sec^−1^ (Supp Table 1). Accordingly, a mean inactive period of ∼ 0.8 sec [= (1/2.5) + (1/2.5)] is measured for single dynein (Fig 2C and related discussion). The predicted mean inactive period for a dynein-team (*N* ≥ 2) is calculated for different models (main text). The best agreement is seen for the Entropic Constraint model using an interaction parameter of Φ = 4.54 (for LPs) and Φ = 2.5 (for SSLBs). Bars are SEM. B) Splitting probability *p(m,N)* for *N* = 5 superstalled dyneins to detach with *m* = 0 to 5 dyneins in the active state (***D***) versus (*N – m*) = 5 to 0 dyneins in inactive state (***D\****). The inset shows a representative stall with different time points marked: *T_0_* is when superstall was initiated, *T_1_* is end of superstall and cargo detachment to fall back to the trap centre, and *T_2_* is end of the inactive period (= *T_2_* – *T_1_*), after which dynein again attaches to the MT to generate force. C) A subset of data (only *N* = 4,5,6 dyneins) from Fig 2E for inactive periods of LPs after Superstalls is re-plotted as a cumulative histogram. Red line shows the cumulative histogram predicted by the Entropic Constraint model using (Φ = 4.54) to represent the superstall condition (all dyneins forced into ***D\**** state). The time constants obtained by fitting experimental data to a two-stage hypo-exponential process are shown. Time constant predicted by the model are also shown (see main text). D) Trajectories of endosome motion obtained from *in-vitro* motility assays with Tug-of-wars (ToWs; start and end shown using red arrowheads) occurring between reversals. <u>Upper Panel</u> shows Plus-to-minus reversals <u>Lower Panel</u> shows Minus-to-plus reversals. MT orientation is shown on the right. E) Duration of ToW (Mean ± SEM) for Minus-to-plus reversals and Plus-to-minus reversals.

Equation (1) describes the mean inactive period starting from the all-inactive state, where the first term corresponds to any of the *N* dyneins becoming active and the second term is the MT-binding rate of this active dynein. This “No-interaction” model predicts significantly faster activation, deviating clearly from the experimentally observed inactive periods that increase steadily with *N* (Fig 4A). Therefore, surprisingly, dyneins within a team must be interacting in some way to delay each other from generating force. This delay amplifies as the size of the dynein team (= *N*) increases to cause a paradoxical form of collective inactivity (Fig 4A).

*What might cause such collective inactivity of dynein-teams after they have generated force?* We proposed that detachment from catch-bond state (***D^CB^*** + ***MT*** → ***D***^∗^; see Fig 2F) forces an inactivating deformation (***D\**** state) in single dynein, which can return to active (***D***) state only via internal rearrangements aided by thermal diffusion. If two or more ***D\**** dyneins are in close proximity (i.e. within some entanglement zone), then entropic constraints could prevent thermal diffusion to delay this activation. Put simply, the dyneins could sterically hinder each other’s diffusion back into the active (***D***) state. Such a situation may become more prominent when dyneins cluster together to form a dynein-team on a lipid raft (size ∼100-200 nm) on late phagosomes (18). We therefore developed a simple two-state (*σ* = 0,1) **<u>Entropic Constraint Model</u>** to quantify the effects of such entanglement within a dynein-team. Similar models have interrogated receptor-ligand interactions bathed in a surrounding thermal reservoir (47). Let us assume that a given inactive (***D\****) dynein can exist in one of two states:-

1. **“Free”** (σ = **0**) This ***D\**** dynein has no other ***D\**** dyneins within its entanglement zone, and can therefore make the ***D***^∗^ → ***D*** transition with a “bare” rate *k*_0_ = 2.5 *sec*^−1^. This is the same rate at which a single dynein undergoes the (***D***^∗^ → ***D***) transition, and was obtained by measuring the additional period for which dynein was inactive after catch-bond in a single-molecule assay (Fig 2C). The choice of *k*_0_ = 2.5 *sec*^−1^ is justified because a single catch-bonded dynein has no other ***D\**** dyneins to entangle with.
2. **“Entangled“** (σ = **1**) This ***D\**** dynein has at least one other ***D\**** dynein within its entanglement zone, and is therefore entropically constrained so that it’s (***D***^∗^ → ***D***) transition is slower (rate *k* < *k*_0_; see Fig 2F). Interaction with other ***D\**** dyneins stabilizes the entangled dyneins, leading to a free energy difference Δ*E* = (*E_Entangled_* − *E_Free_*) < 0.

The free energy of a ***D\**** dynein is *E* = [*E*_Free_ + *σ* · (Δ*E*)] and its partition function is 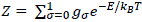. Here *g*_σ_ denotes the degeneracy in state *σ*. Assume a cargo with *N* dyneins capable of generating force, where *m* dyneins (*m* ≤ *N*) are in ***D***, and (*N* − *m*) dyneins are in ***D\**** state. As a simplest model, we allow a given ***D\**** dynein to entangle with only one of the remaining (*N* − *m* − 1) ***D\**** dyneins. Hence, the *σ* = 1 state has a degeneracy of *g*_1_ = (*N* − *m* − 1). The partition function of this system is then

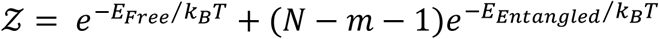

The probability that a given inactive dynein exists in the free state is given by

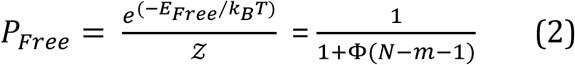

where Ф = ***e***^−Δ***E***/***k***^***^BT^*** is an “entanglement parameter” that measures the entanglement of dyneins in terms of the free energy difference Δ*E*. The value of Φ depends on density and geometrical distribution of dyneins on the cargo membrane, and as discussed later, may change as the endosome/phagosome matures (18). The case Φ = 0 represents a limit (*k* = *k*_0_) where individual ***D\**** dyneins in a team are well-separated and therefore unable to constrain/entangle with each other (Eqn. 1 and grey curve in Fig 4A). If a single ***D\**** dynein has a bare activation rate of *k*_0_ (= 2.5 *sec*^−1^), then the per-dynein activation rate of a ***D\**** dynein in presence of all the other ***D\**** dyneins is

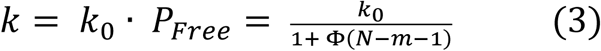

Equation (3) predicts how dynein’s bare activation rate (*k*_0_) is suppressed by other catch-bonded (inactive) dyneins, and will be used next to understand the kinetics of (***D***^∗^ → ***D*** → ***D*** + ***MT***) transitions in an *N*-dynein team leading to collective inactivation.

### ENTROPICALLY CONSTRAINT DIFFUSION: PREDICTION VERSUS EXPERIMENTS

Fig 4B (inset) shows three time-points (*T*_0_, *T*_1_, *T*_2_) associated with the superstall data of a dynein-team on an LP. Based on Supplementary Fig 3A, all dyneins get catch-bonded at superstall (i.e. at *T*_0_). A catch-bonded dynein detaches into the ***D\**** state, from where it may transition to the active ***D*** state during the superstall (i.e. between *T*_0_ and *T*_1_). When the last catch-bonded motor unbinds at *T*_1_, the *N* unbound motors are distributed between ***D*** and ***D\**** states with certain probabilities. We define *p*(*m*, *N*) as a “splitting probability” that the cargo detaches with exactly (*m*) dyneins in active (***D***) state and the remaining (*N* − *m*) in inactive (***D\****) state. The splitting probabilities *p*(*m*, *N*) have been calculated in Appendix-2 (Supplementary Information). Any one of these ***D\**** dyneins can get activated with the modified rate *k* of eqn. (3), yielding an effective rate (*N* − *m*)*k* for the ***D\**** → ***D*** process. Any of the (*m*) active ***D*** dyneins can also undergo ***D*** → ***D*** + ***MT*** transition with rate *π*, yielding an effective rate (*mπ*) to generate force and end the inactive period. With these assumptions, we obtain a set of recursive relations that describe the inactive periods *T*(*m*, *N*) in Appendix-1 (Supplementary Information). The mean inactive period *T*(*N*) predicted by the entropic constraint model can then be calculated using *T*(*m*, *N*) weighted by the splitting probabilities *p*(*m*, *N*).

As shown in Fig 4A, the best agreement of *T*(*N*) with experimental data is obtained for Ф = **4**. **54**. Note that only one unknown parameter (Φ) was varied to find the best agreement. Experimentally measured single-dynein rates (*k*_0_ = 2.5 *sec*^−1^; *π* = 2.5 *sec*^−1^) were used, and values of *N* were inferred from force measurements. From Φ = *e*^−ΔE/kBT^it follows that the entangled state is stabilised by a free energy difference

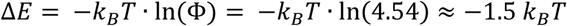

The predicted values of *T*(*N*) are slightly lower than the experimentally observed values for *N* = 5 and 6 (see Fig 4A). This deviation likely follows from our simplifying assumption that any ***D\**** dynein entangles with only one other ***D\**** dynein. As more dyneins cluster together on the LP membrane, a given ***D\**** dynein would encounter multiple entanglements to remain inactive for longer. A more elaborate model allowing for these higher-order effects should agree better with the experiments, but is outside the scope of this manuscript.

We will now test how the value of Φ = 4.54 applies to a specific cargo with *N* ≈ 5 superstalled dyneins. The splitting probability for *N* = 5 case, as shown in Fig 4B, reveals two dominant populations (i) 75.4% with ***D\**** = 5, *m* = 0 and (ii) 20.7% with ***D\**** = 4, *m* = 1. The distribution of inactive periods is therefore dominated by a slow and a fast process relating respectively to the (*m* = 0) and (*m* = 1) starting states. For the complete (***D***^∗^ → ***D*** → ***D*** + ***MT***) transition, the total escape rate of leaving from a state with *m* active dyneins is

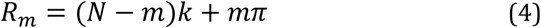

Eqn (4) depicts that each of the (*N* − *m*) dyneins in ***D\**** state can undergo the ***D***^∗^ → ***D*** transition with an entropically constrained rate *k* given by equation (3), while any of the *m* dyneins in ***D*** state can undergo the ***D*** → ***D*** + ***MT*** transition with a rate *π* and generate force. Starting from the *m* = 0 state, because the binding channel is closed, escape is rate-limited by the first activation event which serves as the slowest timescale in the process. Once a single motor is active, the binding channel is open, and escape is faster. This fast timescale governs the *m* = 1 population directly, and also the post-activation phase of the *m* = 0 trajectories. The inactive period distribution is therefore well approximated by a two-timescale form: a slow timescale set by the *m* = 0 first-activation bottleneck and a fast timescale common to both populations. We therefore use the single dynein MT-binding rate (*k*_0_ = 2.5 *sec*^−1^), an entanglement parameter Φ = 4.54 and *N* = 5 in equation (3) to predict the below rates:-

(***State m*** = **0**) … Using equation (3) we obtain *k* = 0.13 *sec*^−1^. Using equation (4) with *m* = 0 yields *R*_0_ = [(5) ∗ (0.13)] = 0.65 *sec*^−1^. The slow timescale governing escape (= 1⁄*R*_0_) is therefore 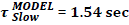.

(***State m*** = **1**) … Using equation (3) we obtain *k* = 0.17 *sec*^−1^. Using equation (4) with *m* = 1 yields *R*_1_ = [(4 ∗ 0.17) + (1 ∗ 2.5)] = 3.18 *sec*^−1^. The fast timescale governing escape (= 1⁄*R*_1_) is therefore 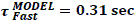.

Recall from Fig 2G that the inactive period after superstalls has *CV* < 1, suggesting a hypo-exponential process involving sequential ***D***^∗^ → ***D*** → ***D*** + ***MT*** transitions. Fig 4C shows the cumulative distribution function (CDF) of inactive periods obtained from Superstalls on Late phagosomes (LPs). The stall forces suggest that these LPs were driven by 4 to 6 dyneins (average *N* ≈ 5 dyneins). Because the splitting probabilities suggest that ***D***^∗^ → ***D*** → ***D*** + ***MT*** transitions are dominated by a slow (*m* = 0) and a fast (*m* = 1) process (Fig 4B), we fit the CDF of inactive periods to a two-stage hypo-exponential:-

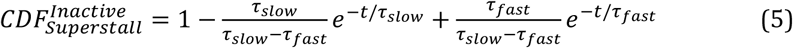

This fit yields experimentally obtained time constants as 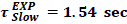 and 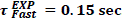. The predicted slow time constant (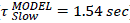) matches the experimental value exactly, but the predicted fast time constant (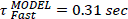) is slightly longer. Note that the full distribution is determined by a cascade of activation rates (*R*_0_, *R*_1_, *R*_2_, *R*_3_, *R*_4_), whereas our approximation assumes only 1⁄*R*_0_ and 1⁄*R*_1_ as the dominant timescales. Using the full distribution hastens exit from inactivity to yield excellent agreement between prediction and experiment (See Figure-1 of Appendix-3 in Supplementary information). Overall, for superstalled cargos, the state with all-inactive dyneins (*m* = 0) at the point of MT-detachment is dominant, and acts as a bottleneck to force generation. For a single dynein, the 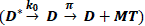 transitions have rates *k*_0_ = 2.5 *sec*^−1^, and *π* = 2.5 *sec*^−1^. For a team of *N* ≈ 5 dyneins forced to detach into ***D\**** state, the (***D***^∗^ → ***D***) transition is slowed down in a manner that can be explained through the Entropic Constraint model of equation (3).

### RELEVANCE OF DYNEIN’S CATCH-BOND TO BIDIRECTIONAL MOTION

The application of a superstall force Δ*F* = (0.3 × *F*_STALL_) over and above the force *F*_STALL_ that dyneins can themselves generate allowed us to “prepare” an *N*-dynein team with all/most dyneins in catch-bonded ***D\**** state. These data led to the Entropic Constraint Model. However, one may ask whether ToWs actually cause dyneins to catch-bond on bidirectionally moving cargoes? Fig 4D shows the video tracks of single endosomes purified from *Dictyostelium* as they moved along a MT *in-vitro*. Endosomes reverse direction frequently as they move (6), exhibiting low-velocity regions where they visibly elongate along the MT (ToWs; red arrows in Fig 4D). Many ToWs exceeded several seconds, with the average ToW duration for Plus-to-Minus or Minus-to-Plus reversals being ∼1.3 seconds (Fig 4E). These ToWs involve 1-2 kinesins against a team of 5-6 dyneins (6). The average *T_STALL_* for beads driven by ∼5 dyneins is 0.75 sec (22). Events with *^T^_STALL_* ≥0.75 sec have a longer inactive period after the stall (Supplementary Fig 2B), suggesting that the optical trap induces catch-bonding in these dyneins. Because the ToW on endosomes often exceeds 0.75 sec (Figs 4D, 4E), a significant fraction of dyneins on endosomes should get catch-bonded during ToWs against kinesin. These dyneins likely enter the inactive (***D\****) state when they detach, making the Entropic Constraint Model relevant to ToWs on endosomes/phagosomes, a claim also supported by the increased superstall-induced reversals of early phagosomes (Figs 3B and 3D).

### DYNEIN ON ENDOSOMES & PHAGOSOMES: TO CLUSTER OR NOT TO CLUSTER

Recall that the characteristic bidirectional motion of early endosomes/phagosomes could not be replicated in MC simulations (Figs 1D, 1E). This discrepancy seeded the subject of this study, namely a possible inactive state in catch-bonded dyneins. We therefore returned to the MC simulation of *N*_+_ = 1 or 2 kinesins and *N*_–_= 6 dyneins to understand how dynein’s inactivity impacts intracellular motion and consequent maturation of endosomes/phagosomes. Dyneins were now kept inactive in the simulation once they had detached from catch-bonded state after a ToW, and were allowed to generate force only after completing the (***D***^∗^ → ***D*** → ***D*** + ***MT***) transitions with kinetics derived from the entropic constraint model. We had obtained a value of Φ = 4.54 by fitting the mean inactive period of LPs to the entropic constraint model (Fig 4A). However, using Φ = 4.54 in MC simulations yielded a high incidence of ToWs with net motion biased in plus direction (Supplementary Figs 4B and 4C). This happens because the high value of Φ keeps dyneins inactive for long periods after every ToW with kinesins. *How then can LP motion be dominated by dynein inside cells with very few incidences of ToW* (18), as also shown in Supplementary Fig 4D. Dyneins use the ORP1L-Rab7-RILP pathway to bind cholesterol on membranes (48), thus clustering into cholesterol-rich lipid rafts to form large dynein-teams. Kinesin has no known preference for cholesterol on these cargoes, and is therefore likely excluded from dynein-rich lipid rafts. This segregation prevents ToWs, so that dyneins rarely get catch-bonded. The entropic constraint model (and Φ) therefore become irrelevant, ensuring that large dynein-teams remain active on late endosomes/phagosomes to drive them rapidly towards lysosomes for degradation.

EPs and LPs of same size have approximately the same number of dyneins, but EPs contain significantly less cholesterol than LPs. Dynein therefore cannot cluster, and is distributed uniformly to get diluted out across the EP membrane (18). This arrangement must “mix up” dyneins and kinesins, and must also reduce dynein-to-dynein proximity to lower Φ on the EP membrane. Do EPs actually have a lower Φ than LPs (Φ = 4.54)? One may answer this question by measuring the mean inactive period between ***DD*** pair-events after superstalling EPs (experiments similar to LPs in Fig 4A). However, superstalling dynein on EPs caused ∼80% of EPs to reverse direction (see Fig 3B), and thus yielded ***DK*** pair-events rather than ***DD*** pair-events. To avoid this problem, we used spherical supported lipid bilayer cargoes (SSLBs; described earlier) where dynein can be recruited in the absence of cholesterol, and additionally the SSLBs do not reverse direction in the trap (37).

Superstall experiments on SSLBs yielded mean inactive periods consistent with lower entropic constraints on dyneins (Φ = 2.5; Fig 4A). Fig 5A shows simulated trajectories of a cargo for Φ = 2.5. Fig 5B shows the fraction of time spent by a simulated cargo in different motility states, along with the same fractions calculated for endosomes from tracks such as in Figs 1A and 4D. The simulated motion with Φ = 2.5 closely resembles the bidirectional motion of early endosomes/phagosomes. Further, simulations showed ∼80% probability that between 3-5 dyneins will get catch-bonded during ToWs, and will therefore enter an inactive (***D\****) state after ToW (Fig 5C). Thus, the entanglement/constraint effect indeed appears lowered when dyneins are not clustered together into cholesterol-rich domains. The result is that dispersed dyneins form smaller teams to now compete in ToWs against kinesins and generate bidirectional motion of EPs/early endosomes (18).

**FIGURE 5.**
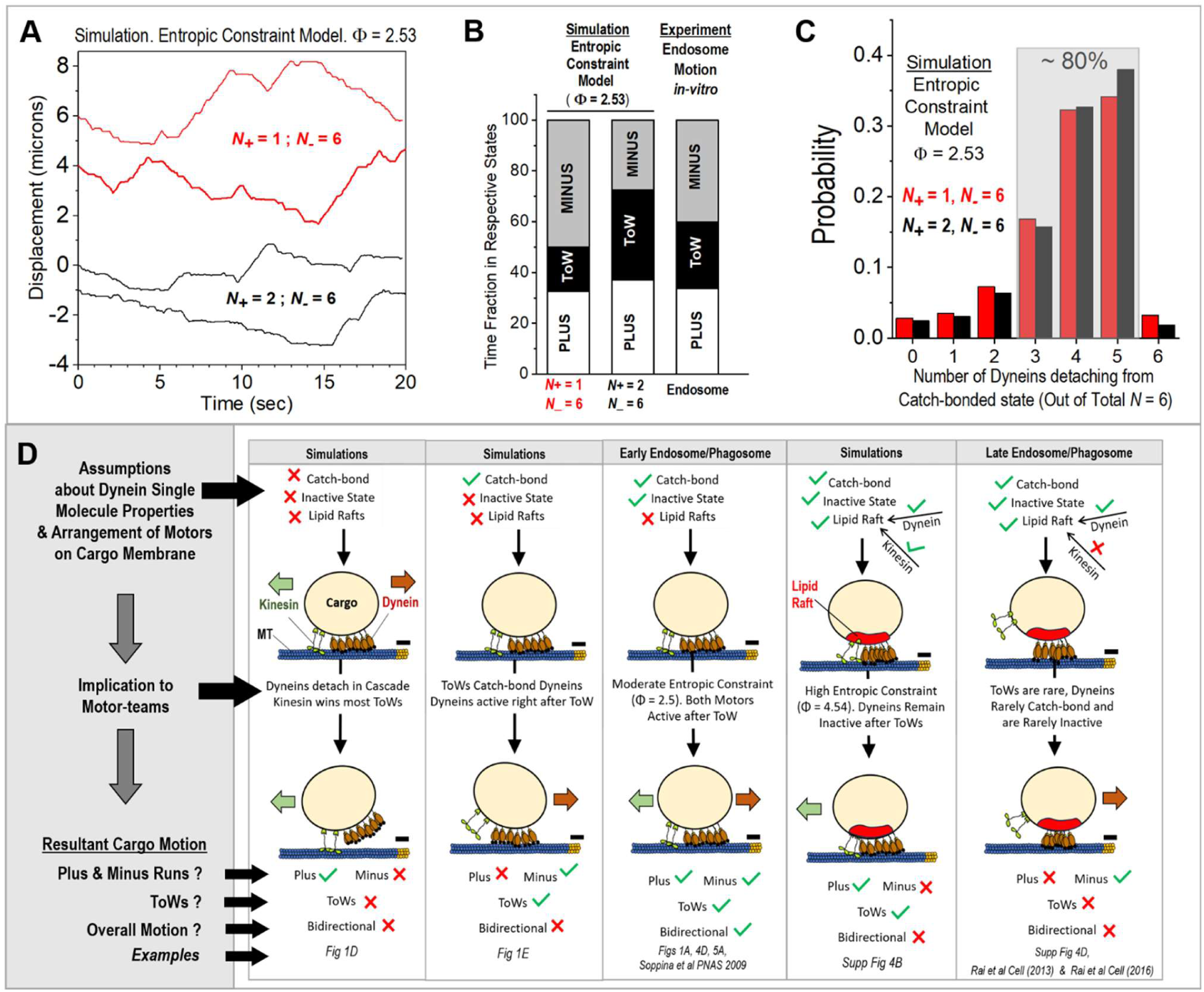
A) MC simulation of cargo motion for the indicated total numbers of kinesin (*N*_+_) or dynein (*N*_–_) motors. These simulations use *k_o_* = 2.5 s^−1^ and Φ = 2.53 and are based on the entropic constraint model (equation 3). B) The time fraction spent in different motility states (Plus, ToW or Paused, Minus) obtained from simulated motion for *n* =1000 trajectories of cargo motion driven by the mentioned number of kinesin (*N*_+_) or dynein (*N*_–_) motors (see Fig 5A). See the Methods section for details related to segmentation of simulated tracks into different motility states. The respective time fractions for experimental data of purified endosomes moving *in-vitro* along MTs is also shown for comparison (video tracked data from *n* = 56 endosomes used, see figures 1A, 4D). C) The probability that a certain number of dyneins (indicated along *X*-axis) will get catch-bonded, and therefore detach from the MT into an inactive (***D\****) state after a ToW against kinesin. These probabilities are calculated from the simulations that are shown in Fig 5A. Note that a large fraction (∼80%) of dyneins is in ***D\**** state after ToW-induced detachment. D) Possible scenarios for ToW-mediated motion of cargoes driven by a few (1 to 2) Kinesins and ∼6 Dyneins. Top row depicts the different situations, where a single dynein motor is allowed (**√**) or not allowed (**×**) to enter into an inactive (***D\****) state after Catch-bond formation. The top row also depicts whether a cargo is allowed to have Lipid Rafts. If Lipid Rafts are allowed, then are Kinesin and Dynein allowed into these Lipid Rafts? The expected events occurring on cargoes are shown in the middle row. The bottom row shows the net output, i.e. the nature of cargo motion under different scenarios. As indicated, only two of these scenarios are observed inside cells (Early Endosome/Phagosome and Late Endosome/Phagosome). For more details, see main text and Discussion.

## DISCUSSION

Cells must continuously generate endosomes/phagosomes that mature through “kiss-and-run” interactions with other vesicles. This maturation is impossible unless endosomes/phagosomes explore cellular space by moving in back-and-forth (bidirectional) manner along MTs (16, 46). ToW-mediated bidirectional motion of endosomes is therefore critical for cells to absorb nutrients via endocytosis and to recycle endocytic receptors. Similarly, bidirectional motion of phagosomes is essential for immune cells to degrade pathogens (16, 18). Cytoplasmic dynein is a central player in bidirectional motion, competing against many kinds of kinesins and perhaps also against other impediments that motile cargoes face inside cells (1, 3). Dynein may achieve such diverse functions by virtue of its complex architecture, the many adaptors that it binds and its ability to work in a team (31). While some *in-vitro* bead experiments measure higher force (49, 50), there is significant evidence that dynein is a weak motor generating between 1 to 2 pN force on native-like purified phagosomes or phagosomes and lipid droplets moving inside cells (18, 22, 23, 51, 52). Multiple dyneins must therefore often form teams to resist stronger kinesins during ToWs (AMCM; Fig 1B).

Cargo motion may depend on how shear force experienced during ToW affects the dynein-MT bond. Receptor-ligand interactions usually display slip-bond behaviour where the detachment rate increases exponentially with opposing force (44), as shown in Fig 1C. If dyneins within a team were to slip-bond during ToW, then they would detach rapidly in a cascade (31, 53). ToWs could never be sustained, and kinesins would always dominate (see Fig 1D). This explains why dynein requires a catch-bond if it has to work in a team. However, dynein’s catch-bond creates the opposite problem. Cargos can move freely in minus direction, but the dyneins catch-bond as soon as they sense force from kinesin, causing all motion to get blocked (Fig 1E). *How intracellular vesicles can <u>actually</u> exhibit bidirectional motion therefore remains unclear.* Based on the evidence found here, a possible resolution to this question will be discussed below through *three effects*:-

### 1) Dynein’s Asymmetric Response to Force

Opposing force strengthens the dynein-MT bond, i.e. it induces a catch/slip-ideal bond response (54) in a manner that likely prevents cascaded detachments within a dynein-team (53). Contrarily, assisting force weakens the dynein-MT bond to detach it faster, i.e. it induces a slip-bond response possibly minimizing any drag that bound dyneins could exert on their motile partners (54). The structural basis of this asymmetric response is not fully understood. It may derive from force-induced changes in dynein’s coiled-coil stalk, from changes directly in the MT binding domain, or from strain-induced changes in the MT lattice (54). Dynein’s stalk binds to the MT lattice through multiple ionic interactions (salt bridges, hydrogen bonds) that are highly sensitive to orientation (55). These interactions switch between *<u>intra</u>*molecular versus *<u>inter</u>*molecular partners, respectively stabilizing low or high affinity binding states of dynein to MTs as the motor goes through its mechanochemical cycle. Substituting negatively charged interfacial residues (CC1-E3289 and H6-E3378) with hydrophobic alanines actually increases dynein’s affinity to MTs, strengthening the argument that dynein has evolved to function with submaximal processivity (55).

### 2) Inactivating Deformations in Single Dynein when it Detaches from Catch-bond

The ***D^CB^*** + ***MT*** state (Fig 2F) displays a catch/slip-ideal bond response likely because its orientation favours high-affinity intermolecular interactions. This bond eventually breaks against high force (***D^CB^*** + ***MT*** ◊ ***D\****; Fig 2F). *We propose that the aforesaid ionic bonds get reoriented/deformed **<u>during the breakage</u>** to now favour intramolecular partners*. The new orientation therefore induces an inactive state of dynein with low affinity to MTs which persists in the detached ***D\**** dynein. The catch-bonding transition (***D*** + ***MT*** ◊ ***D^CB^ + MT***; Fig 2F) is rapid because it is fueled by force/energy from kinesin or from the trap. However, after dynein has detached (***D^CB^*** + ***MT*** ◊ ***D\****), this force is gone and the following (***D\**** ◊ ***D***) transition can only happen via diffusion. A single ***D\**** dynein must therefore undergo slow thermal diffusion on a timescale of ∼0.4 sec 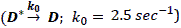 to return to its native conformation before it can again generate force ***(D*** → ***D*** + ***MT)***. The stepping of motors was recently found to cause localized damage/deformation of the MT-lattice. Because a force-induced inactivity is seen here only for dynein (and not for kinesin), the effects observed in our study derive from changes in dynein and mot on the MTs lattice.

### 3) Collective inactivity Enforced by Entropic Constraints: Autoregulation of Dynein-teams

Larger dynein-teams enable longer minus-directed runs (6, 25), and must therefore be kept inactive for longer after a ToW if a comparable extent of plus-directed motion is to be achieved by a few kinesins. Such *collective* inactivity is exactly what we observe through superstall experiments where the mean inactive period increases with dynein number (Fig 4A). For a dynein-team that has detached against high force, multiple ***D\**** dyneins must explore phase space by diffusing around within a confined volume. This process likely gets entropically constrained when many dyneins cluster into a nanoscale lipid-raft domain to form a team [e.g. on LPs; see Ref. (18)]. The entropic constraint grows as more dyneins cluster together, and is the simplest explanation for why inactivity amplifies with dynein number *N* (collective inactivity; Fig 4A). While clustering packs many dyneins together to generate large persistent forces, it also keeps a larger dynein-team inactive for longer after it has generated force. This form of auto-regulation may prevent a “runaway-dynein” situation by ensuring that other force-generating entities (e.g. kinesins) can get a chance during the inactivity of dyneins. The above-described ideas are summarized as a schematic (Fig 5D), and follow from the Entropic Constraint model where the single-molecule rate (*k*_0_) for dynein’s reactivation is replaced by a slower rate [*k*; see equation 3] that arises due to entanglement between inactive (***D***^∗^) dyneins. The Entropic Constraint model causes ∼16-fold delay (= *k*_0_⁄*k*) in the (***D***^∗^ → ***D***) transition for motion driven by *N*∼5 dyneins, which manifests as collective inactivity of these dyneins and allows kinesin to take over during this inactivity.

The proposed entropic effects are quantified here through an Entropic Constraint Model (equation 3). Using a moderate value of the Entanglement parameter (Φ ≈ 2.5) in this model generates an optimal balance of activity/inactivity in dynein-teams, resulting in vigorous bidirectional motion similar to endosomes (Figs 5A, 5B). The same processes likely generates bidirectional motion of early phagosomes (EPs) in immune cells, causing EPs to acquire cholesterol and mature into late phagosomes (LPs). Cholesterol helps to form lipid rafts on LPs, where many dyneins cluster together to elevate Φ (= 4.54) and propel LPs rapidly towards lysosomes. However, this strategy works only if the kinesins have been removed from the lipid rafts to prevent ToWs that would otherwise inactivate the clustered dyneins for long periods (Fig 5D). Our findings could have broader relevance to the mechanobiology of catch bonded systems. For example, the detection of sound waves requires forced bending of tip links in the inner ear, where cadherin forms a catch-bond (56). The dynamic response of tip links should be sensitive to the frequency of sound, and may involve repeated formation/breakage of catch-bonds. If cadherin’s catch-bond also gets inactivated transiently like dynein, this could have interesting implications for the perception of sound at different frequencies.

## MATERIALS AND METHODS

### Further Details of Experimental Procedures, Methods and Reagents Provided in Supplementary Information

J774 (ATCC-TIB-67), HeLa (ATCC-CCL2) and *Dictyostelium discoideum* (AX2-DBS0235521) cells were used as described (6). J774 cells were cultured in DMEM supplemented with 10% heat-inactivated FBS. Cells were grown in a humidified incubator at 37°C and 5% CO_2_. *Dictyostelium* cells cultured in HL5 medium at 22°C were used for phagosome and ATP releasate preparations as described (6). Live-cell imaging and *in vitro* motility assay of EPs and endosomes were performed as described (6). Free or no-load motion of these cargoes was observed using a custom-modified Nikon TE2000 microscope under differential contrast imaging mode (Ref Barak). Optical trap stiffness (*K_TRAP_*) was determined for *in-vitro* assays using the power spectrum method or Video matching method as appropriate (6, 57). *K_TRAP_* values between 0.01 pN/nm to 0.08 pN/nm were measured depending on the laser power used. For *in-vitro* superstall experiments, we rapidly moved the MT-bound cargo away from the trap centre in same direction as the moving cargo using a piezo-controlled stage (6). For *in vivo* stall/superstall experiments, latex beads (750nm diameter) were phagocytosed into J774 cells. Motile EPs were trapped using a custom developed LabView program (25). AVI-format videos of these experiments were analyzed to verify that the trap aligned with the cargo’s free motion before and after trapping. Further details of video tracking/analysis and the Quadrant photo detector setup can be found elsewhere (6, 57). Monte Carlo simulations were implemented in Python and uploaded to GitHub (https://github.com/scipy-aniket/CargoSim). ImageJ was used to post-process the deep-SIM image of the J774 cell. Origin-2021 was used to prepare the graphs.

## STATISTICAL TESTS

Based on the normality of the dataset, a two-tailed *t*-test or a two-tailed Mann-Whitney test was used to assess significance. The statistical relevance was tested for some non-normal data sets using a bootstrapping procedure described earlier (27).

## Supporting information

SuppFigs,Table,Methods,VideoCaptions

## ACKNOWLEDGEMENTS

We acknowledge funding from IIT Bombay (RD/0520-IRCCSH0-010) and Senior Fellowships from the Department of Biotechnology-Wellcome Trust India Alliance (Grants IA/S/11/2500255 and IA/S/19/2/504634). D.C. acknowledges funding from IIT Bombay (RI/0720-10001928-001). We thank the Imaging Facility of IIT Bombay for DeepSIM imaging. We thank Pankaj Kumar, Ashwin D’Souza and Manish Kumar Singh for help with experiments.

## Notes

### Competing Interest Statement

The authors have declared no competing interest.

