## Supplementary material for "Force Induces a Collective Inactivity in Dyneins to Auto-Regulate Bidirectional Transport of Vesicles": SuppFigs,Table,Methods,VideoCaptions

*Dwiteeya Chaudhury et. al.*

Supplementary Information Contains :-

1. Supplementary Figures 1 to 4
2. Supplementary Table 1
3. Supplementary Materials and Methods
4. Supplementary Video Captions (for Videos S1, S2, S3, S4)
5. Entropic Constraint Model (Theory)

   Appendix 1 Mean Inactive Times
   Appendix 2 Splitting Probabilities
   Appendix 3 Distribution of Inactive Times
   (Includes 1 Figure :- *Appendix-3 Figure 1*)

6. References in Supplementary Information

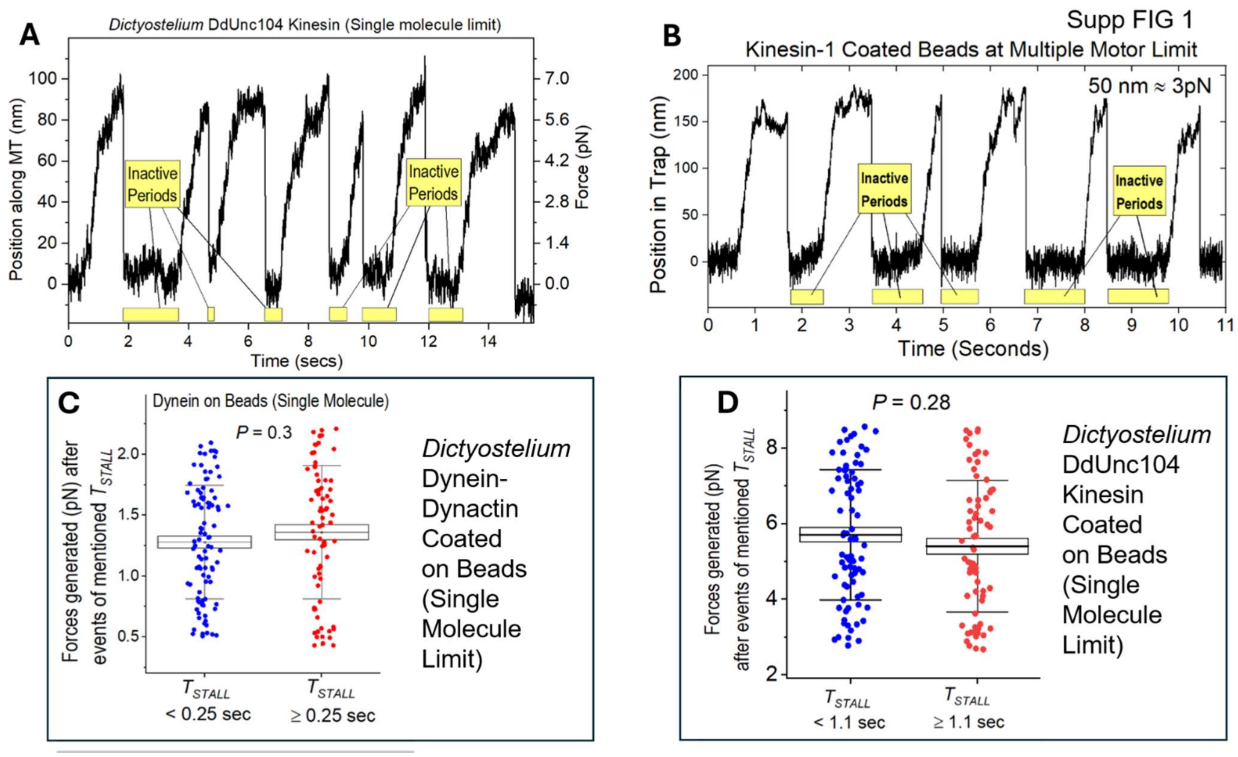


### Supplementary Figure 1

**Long inactive periods are seen between force generation events by Kinesin.**

A) Successive stalls of DdUnc104 Kinesin purified from *Dictyostelium*. The motor was coated on beads at a dilution where <30% of the beads generated force (single molecule limit). The displacement along MT from centre of the trap (left axis) and corresponding force (right axis) are shown. Inactive periods between two successive stalls are indicated. Inactive period is the time taken by the motor to rebind the MT after it has detached from the MT against force from the trap. An inactive period is when the bead is within (3 × SD) distance from the trap centre between two successive force-generation events. SD is the standard deviation in position of a bead trapped away from a MT (no force generation by motors).

B) Same as in (A), but for kinesin-1 purified from goat brain and coated onto beads. This experiment was done at a concentration of motors where ~60% of beads generated force (multiple motor limit). These stalls correspond to force generation expected by two kinesin-1 motors. Inactive periods between successive stalls are indicated.

A) Stalls generated by single dynein are divided into two categories :- Stalls with ($T_{STALL}$ < 0.25 sec) or ($T_{STALL}$ ≥ 0.25 sec). The force generated by dynein is plotted for the **subsequent** event following an event in the aforesaid categories of $T_{STALL}$. Boxes are SEM and whiskers are SD. Mean is also indicated.

B) Stalls generated by single DdUnc104 Kiinesin are divided into two categories :- Stalls with ($T_{STALL}$ < 1.1 sec) or ($T_{STALL}$ ≥ 1.1 sec). The force generated by dynein is plotted for the **subsequent** event following an event in the aforesaid categories of $T_{STALL}$. Boxes are SEM and whiskers are SD. Mean is also indicated.

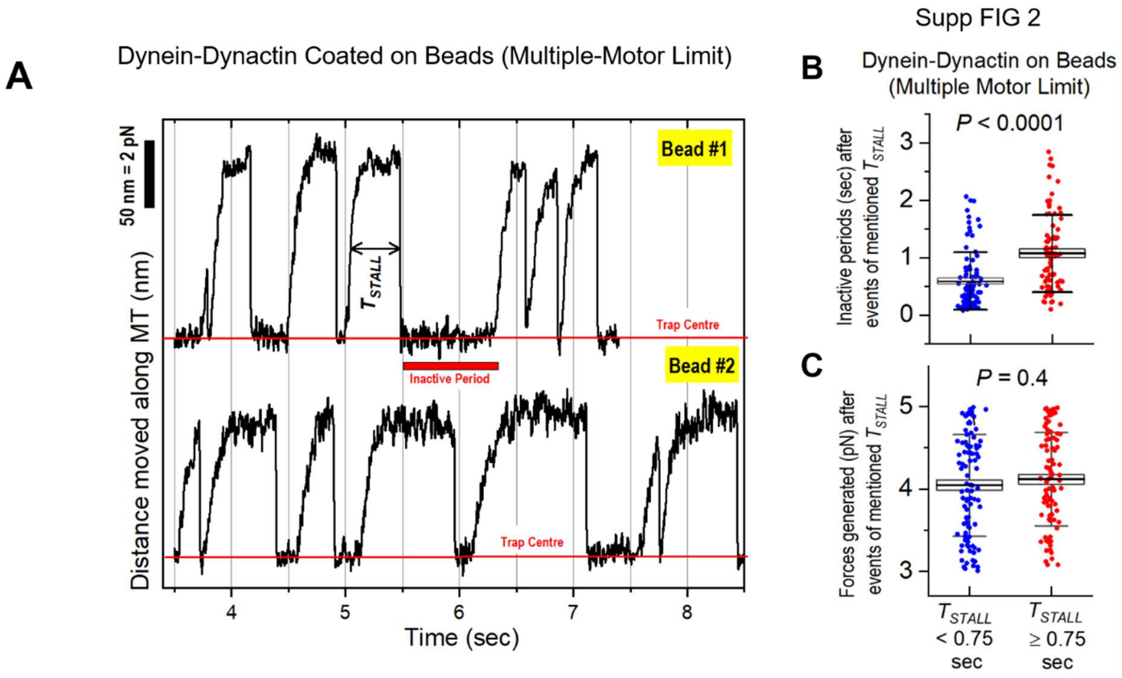


#### Supplementary Figure 2

**Stall durations, Inactive periods and Stall force for dynein coated on beads in the multiple-motor limit.**

C) Motion of two dynein-dynactin coated beads (Bead #1 and Bead #2) against an optical trap at multiple-motor limit. Based on the maximum force generated, these beads are driven by ~4 to 6 dynein motors. Red line indicates the trap centre. The stall duration ($T_{STALL}$) is indicated for one stall. The inactive period after that stall is also indicated. An inactive period is when the bead is within (3 × SD) distance from the trap centre between two successive force-generation events. SD is the standard deviation in position of a bead trapped away from a MT (no force generation by motors).

D) The inactive periods measured through experiments shown in panel A are divided into two categories :- Inactive period after stalls of short duration ($T_{STALL}$ < 0.75 sec) or of a long duration ($T_{STALL} \geq$ 0.75 sec). See main text for the choice of this threshold. The average inactive period is significantly longer after a stall of ($T_{STALL} \geq$ 0.75 sec) category. Boxes are SEM and whiskers are SD. Mean is also indicated.

E) The force generated by dyneins in the force-generating event that occurred after a short stall ($T_{STALL}$ < 0.75 sec) or after long stall ($T_{STALL} \geq$ 0.75 sec). Boxes are SEM and whiskers are SD. Mean is also indicated.

Supp FIG 3

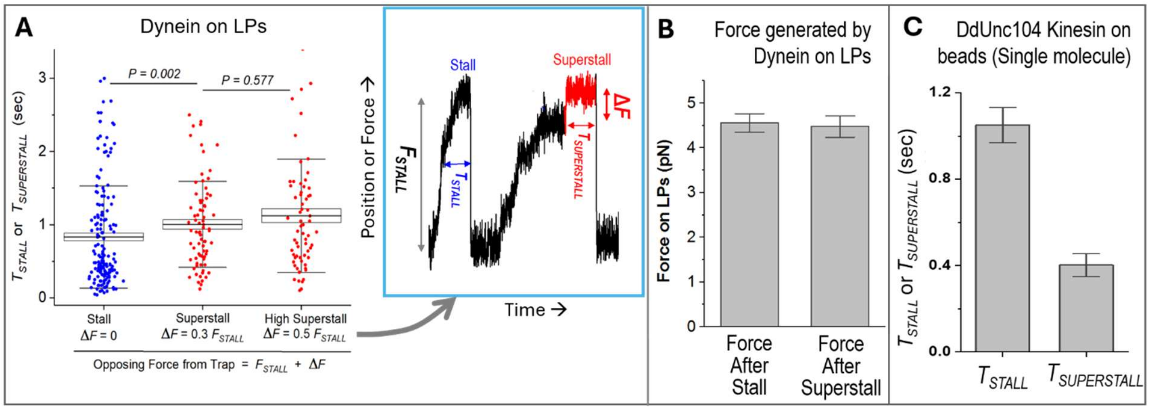


### Supplementary Figure 3

**Dynein-MT and kinesin-MT interactions respectively show catch and slip-bond behaviours.**

A) The time ($T_{STALL}$ or $T_{SUPERSTALL}$) for which dyneins on a Late Phagosome (LP) resist force from an optical trap. ***INSET*** inside blue rectangle explains the meaning of various terms ($T_{STALL}$ ; $T_{SUPERSTALL}$ ; $F_{STALL}$ ; $\Delta F$). $T_{STALL}$ or $T_{SUPERSTALL}$ are plotted along Y-axis under different conditions ***(i) Stall*** Piezo Stage not moved ($\Delta F = 0$) and dyneins generate force $F_{STALL}$ ***(ii) Superstall*** Piezo stage moved to apply additional force $\Delta F = 0.3\times F_{STALL}$ over and above $F_{STALL}$ on dyneins ***(iii) High-Superstall*** Piezo stage moved to apply high additional force $\Delta F = 0.5\times F_{STALL}$ over and above $F_{STALL}$ on dyneins. The $T_{STALL}$ of dyneins ($n$ = 138 stalls measured) is significantly lower than their $T_{SUPERSTALL}$ ($n$ = 76 superstalls measured). This suggests that superstalls induce catch-bond formation in dyneins. $T_{SUPERSTALL}$ is not significantly different between *Superstall* and *High-Superstall* condition ($n$ = 66 superstalls measured), suggesting that the Superstall force ($\Delta F = 0.3\times F_{STALL}$) is sufficient to catch-bond almost all of the dyneins. Boxes are SEM and whiskers are SD. Mean is also indicated. $P$-values are derived from Mann-Whitney tests. The statistical inferences were also verified using a boot-strapping approach.

B) The force generated by dyneins on Late phagosomes after they had been stalled ($n$ = 138 stalls measured) is similar to force generated after they had been superstalled ($n$ = 76 superstalls measured). Error bars are SEM.

C) The $T_{STALL}$ of kinesin ($n$ = 64 stalls measured) is significantly higher than their $T_{SUPERSTALL}$ ($n$ = 64 superstalls measured). This suggests that superstalls induce slip-bond formation in kinesin. Error bars are SEM.

Supp FIG 4

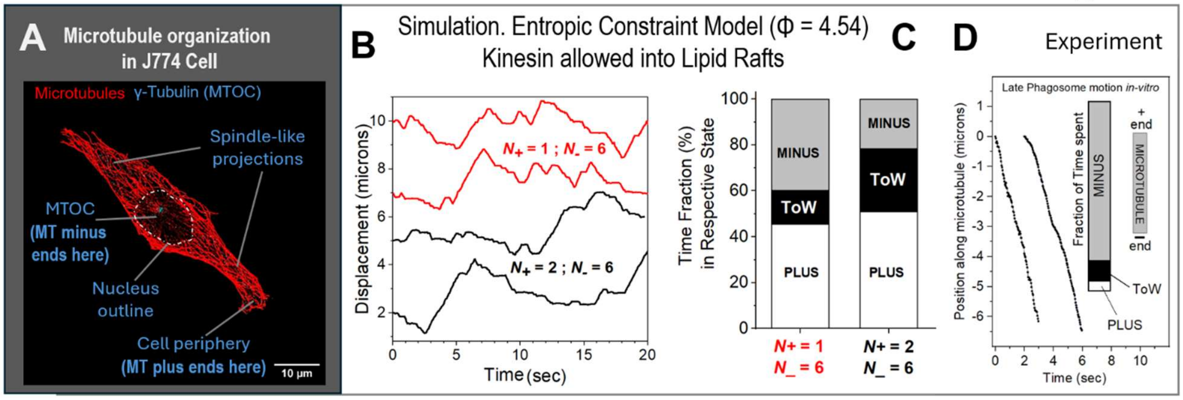


#### Supplementary Figure 4

**MT orientation in J774 A.1 cells and MC Simulation of cargo motion under different parameter regimes of the Entropic Constraint Model.**

A) Structured Illumination Microscope (SIM) image of a J774 A.1 cell stained with anti-α tubulin (red, Alexaflour 568) and anti-γ tubulin (blue, Alexaflour 405). MTs emanate from the MT organizing centre (marked by γ tubulin) and extend in close-to-parallel orientation into the spindle-like regions of the cell.

B) MC simulation of Cargo displacement with mentioned numbers of Kinesin (N+) and Dynein (N-) motors, where both motors are allowed to be proximal to each other on the cargo membrane (e.g. both allowed into lipid rafts). The simulation was done under the entropic constraint model (Equation 3) with interaction parameter $\Phi = 4.54$.

C) The time spent in different motility states (Plus, Paused, Minus) is shown during the simulated trajectories of panel B. Paused state has velocity < 30 nm/sec.

D) Representative motion of late phagosomes (LPs) purified from *Dictyostelium* moving along microtubules *in-vitro*. Microtubule polarity is indicated. Note the dominant minus-directed motion at high velocity driven by dynein-teams. Thus, the MC simulations (panel B and C) **do not** replicate the predominantly minus-directed motion that is known experimentally for LPs. This can be explained if Kinesin is not allowed to be present in close proximity to dyneins on the LP membrane, thereby avoiding ToWs and making Φ irrelevant (see main text).

#### SUPPLEMENTARY TABLE 1

Parameters used for the Monte Carlo simulations and for the interpretation of Experimental data

| Parameter Name | Motor Type and Value of Parameter | Source<br>Note : DOI link is provided specifically for citations that were not included in the Main manuscript Reference list |
|---|---|---|
| No-Load Attachment Rate ($\pi$) | Kinesin-1<br>5 $sec^{-1}$ (Free motion)<br>1 $sec^{-1}$ (In Trap) | Narayanareddy et al Commun. Biol. (2024).<br>Kerssemakers et al PNAS (2006). |
| | DdUnc104 Kinesin | See Fig 2A, Same as Kinesin-1 |
| | Dynein<br>1.6 $sec^{-1}$ and<br>2.5 $sec^{-1}$ | Muller et al PNAS (2008), Gross et al JCB (2002), DSouza et al Nat. Comm. (2023).<br>Also See Figs 2A & 2C |
| No-Load Detachment Rate ($\epsilon_0$) | Kinesin-1<br>1 $sec^{-1}$ | Schnitzer et al *Nat Cell Biol* **2**, 718–723 (2000). https://doi.org/10.1038/35036345 |
| | DdUnc104 Kinesin<br>0.3 $sec^{-1}$ | Calculated from Velocity and Run length data in Soppina et al PNAS (2009). |
| | Dynein<br>0.67 $sec^{-1}$ | King et al *Nat Cell Biol* **2**, 20–24 (2000). https://doi.org/10.1038/71338<br>Muller et al PNAS (2008). |
| Detachment Rate Under Load | Kinesin-1<br>1 to 3 $sec^{-1}$ | Kunwar et. al. PNAS (2011). |
| | DdUnc104 Kinesin | Assumed same as Kinesin-1 based on Soppina et al PNAS (2009). |
| | Dynein | See Fig 1C,<br>also see Kunwar et. al. PNAS (2011). |
| Stall Force ($F_{STALL}$) | Kinesin-1<br>5.5 pN | Barak et al Nat. Methods (2013).<br>Schnitzer et al *Nat Cell Biol* **2**, 718–723 (2000).<br>https://doi.org/10.1038/35036345 |
| | DdUnc104 Kinesin<br>5.5 pN | Soppina et al PNAS (2009). |
| | Dynein<br>1.1 pN | McKenney et al Cell (2010), Nicholas et al Nat. Comm. (2015), Rai et al Cell (2013), Rai et al Cell (2016), Reddy et al Nat. Comm. (2016), Sanghavi et al PNAS (2021), Sanghavi et al Curr. Biol. (2018) |
| Rest length of Motor (Used for MC simulations with) unequal load sharing) | Kinesin-1<br>60 nm | Kerssemakers et al (2006). |
| | DdUnc104 Kinesin | Assumed same as Kinesin-1 |
| | Dynein<br>70-100 nm | Roberts et. al. *Nat Rev Mol Cell Biol* **14**, 713–726 (2013).<br>https://doi.org/10.1038/nrm3667<br>Rai et al Cell (2016) |

### SUPPLEMENTARY MATERIALS AND METHODS

#### MATERIALS

All chemicals were procured from Merck unless otherwise mentioned. HL5 medium was obtained from Formedium. DMEM, FBS, NaCl and Tween 20 were obtained from HiMedia. Pepstatin A was purchased from MP Biomedicals. Penstrep antibiotic was purchased from GIBCO. Both the primary antibodies (AB 11317 – γ-tubulin and T6199 – α-tubulin) used for immunostaining were a kind gift from the Swapnil Shinde Lab. Secondary antibodies were obtained from Invitrogen. The polystyrene beads used in all the experiments were 750nm in diameter and procured from Polysciences. Strep-Magnetic bead was purchased from Magcellect. BSA was purchased from GeNei.

#### METHODS

##### Cell Culture

J774 (ATCC-TIB-67), HeLa (ATCC-CCL2) and *Dictyostelium discoideum* (AX2-DBS0235521) cells were used. J774 cells were cultured in DMEM supplemented with 10% heat-inactivated FBS. The cells were passaged at 70-80% confluency using a rubber policeman. HeLa cells were cultured in DMEM with 10% FBS and passaged at 90% confluency with trypsin-EDTA. Cells were grown in a humidified incubator at 37°C and 5% $CO_2$.

*Dictyostelium discoideum* cells were cultured in HL5 media at 22°C. These cells were maintained in petri plates and passaged at 90% confluency. For phagosome and motor protein preparation, these cells were cultured in suspension by shaking at 165 rpm.

##### Early Phagosome and Endosome Motion Inside Cells

J774 cells were seeded into a custom-built glass-bottomed Petri plate to a final confluency of 40%. These cells were grown for 24 hrs in slightly acidic complete media (pH 7.2), allowing them to spread and form a spindle shape. Polystyrene beads were washed thrice and then diluted (1:50) in incomplete DMEM. 120 µl of these diluted beads was added to the chamber dish and kept at 37°C. After 10 minutes, the medium was changed to wash out excess beads. Then the cells with phagocytosed beads in the chamber dish were imaged in Differential Interference Contrast (DIC) mode using a 100X oil-immersion objective on a Nikon TE2000 microscope. The objective was preheated to maintain 37°C for live-cell imaging of early phagosome motion. Similarly, HeLa cells were also seeded and grown overnight on a custom-built chamber in complete media. Rhodamine-Dextran (2mg/ml) was added to label endosomes within cells for 20 mins. These endosomes were imaged simultaneously in DIC and fluorescence using the same setup mentioned above. Videos of early phagosome and endosome motion were acquired, tracked and analysed as previously described in [1].

##### Immunostaining Protocol and Imaging

J774 cells were seeded on acid-washed coverslips in slightly acidic complete media (pH 7.2) to achieve spindle-shaped cells. After achieving 80% confluency on the coverslips, the cells were fixed using 4% PFA for 15 mins at room temperature. Then a gentle wash was given using 1X PBS. The coverslips were incubated in pre-cooled methanol at -20 °C for 5 min. This step is important for the immunostaining of γ–tubulin. The coverslip was washed twice in 1X PBS to rehydrate the cells after the methanol treatment. Following this, the cells were permeabilised with 0.1% TritonX-100 in PBS for 30 min and blocked with PBST (0.1% Tween20 in PBS) containing 5% BSA for 1 hr at room temperature. After blocking, the cells were incubated with α-tubulin (1:1000 dilution in 5% BSA-PBST) and γ–tubulin (1:500 dilution in 5% BSA-PBST) primary antibodies overnight at 4°C. The next day, the coverslip was carefully washed twice with PBS, incubated with secondary antibodies (1:1000 dilution in 5% BSA-PBST) for 1hr at room temperature in a humid chamber, and mounted for imaging.

These cells were imaged using a Nikon Ti2-E microscope platform with the X-light V3 module in Deep SIM mode and a 100X oil-immersion objective. α-tubulin and γ–tubulin were imaged in the red channel (568nm) and the blue channel (405nm), respectively.

##### Tubulin Isolation

Fresh brain from one goat was homogenised after cleaning and removing blood vessels in 70ml of homogenising buffer (50mM Pipes pH-6.8, 3mM $MgCl_2$, 1mM EGTA, 0.1% β-mercaptoethanol, 1.5x Protease Inhibitor Cocktail (PIC) and 2mM PMSF) using a mixer grinder. The homogenate was centrifuged for 1hr at 19000rpm, 4°C. Supernatant was collected after the spin and transferred to a glass measuring cylinder placed in the water bath at 37°C. For the first polymerisation step, an equal volume of 2 M Glutamate was added to the supernatant, followed by 10% DMSO and 0.5 mM GTP at 37 °C. The mixture was intermittently mixed every 10 mins for an hour, after which it was centrifuged at 19000rpm,30°C for an hour to collect the pellet containing impure polymerised microtubules. Then, the first depolymerisation was performed in 20 mL of depolymerisation buffer (50 mM Pipes, pH 6.8, 3 mM $MgCl_2$, 1 mM EGTA, 0.5x PIC and 0.1 mM GTP), using a Dounce homogeniser on ice for 1 hour. The solution from the Dounce was centrifuged for another hour at 19000rpm, 4°C. Supernatant was collected again, followed by a second polymerisation and depolymerisation step to increase the purity of the collected tubulin. The second depolymerisation step was performed for 2.5-3 hours, and the volume was kept low (~5ml). The protein concentration of the supernatant collected after the second depolymerisation was measured to be 6mg/ml. 5 µL aliquots of tubulin were made and snap-frozen for long-term storage in liquid nitrogen.

**Tubulin Polymerisation**

6mg/ml tubulin was mixed with BRB80 (80 mM PIPES, 1 mM EGTA, 1 mM $MgCl_2$) and 1 mM GTP to maintain the tubulin concentration within the polymerising range (2-3mg/ml). The mixture was polymerised at 37°C for 45 minutes. After polymerisation, 3 µM BT (0.3 µl of 10mM Taxol in 1ml BRB80) was added, and the reaction was diluted to 200 µl using a cut tip. Different dilutions were made based on the efficiency of polymerisation.

**Preparation Of Polarity Labelled Microtubules**

Microtubules were labelled to identify the minus end in DIC microscopy for the in-vitro motility assay, as described in [2]. Briefly, biotinylated tubulin was polymerised in the presence of 60% glycerol, 1 mM GTP, 100 mM $MgCl_2$, and BRB80 at 37°C for 30 minutes. After polymerisation, taxol was gradually added to stabilise the microtubules, ensuring the effective concentration remained below 10 µM. These biotinylated microtubules were sheared 40-60 times to form small seeds (short length microtubule segments) using either a gel-loading tip or an 18-G syringe needle. These seeds were incubated with unlabelled tubulin (1:2 ratio) with 1 mM GTP, and BRB 80 at 37°C for 45 min. This step is known as extension of the biotinylated tubulin seeds. These minus-end labelled microtubules were flowed in motility chambers and were tagged using streptavidin-coated magnetic beads (150nm diameter)

**Isolation Of Phagosomes And Post-Nuclear Supernatant (PNS) From *Dictyostelium* Cells**

1. Late Phagosomes

Cells grown in 100 ml suspension culture were pelleted down at 1000g, 4°C for 5 mins and resuspended in 4.5ml of chilled Sorensen's buffer (pH=6.5). Note that the pH of the Sorensen's buffer should match the HL5 media. Separately, Polystyrene beads were washed three times using HL5 media (10,000 rpm, 5 min) and resuspended in 500 µl of chilled Sorensen's buffer. These beads were added to the cells (total volume = 5ml), and the mixture was incubated for 30 minutes at 4 °C with very slow mixing. This step is called synchronisation. Then, the mixture was shaken at 160 rpm in 100 mL HL5 media at 22°C for 15 min. This step is known as pulse. 330 ml of chilled Sorensen buffer was added into 100 ml cell suspension to stop the pulse. Next, the cells were centrifuged at 1000g for 5 min at 4°C. The pelleted cells were resuspended again in Sorensen buffer (5ml) and added to fresh 100 ml HL5 media (shaking, 22°C) for 45 mins. This step is known as chase. The phagocytosed beads mature into Late Phagosomes during chase. The chase was stopped by adding 330ml of chilled Sorensen's buffer, as mentioned earlier. The cells were centrifuged again at 1000g for 5 min at 4°C. This pellet was washed three times using Sorenson's buffer to remove non-phagocytosed beads. The pellet was weighed and lysed on ice with 1 ml lysis buffer – LB30 (30 mM Tris pH 8, 4 mM EGTA, 30% sucrose) containing all inhibitors (1x PIC, 20µg/ml Pepstatin A, 1.75mg/ml Benzamidine, 6mM DTT, and 1.75mg/ml PMSF) and 1.5 mM ATP for 1 g cell pellet maintaining a 1:1 (w/v) ratio. Lysis was performed using a 5 µm pore filter and a syringe. The cell lysate was gradually layered on top of a 25% sucrose gradient (LB 25: 30 mM Tris pH 8, 4 mM EGTA, 25% sucrose) with all inhibitors and spun in an MLS50 rotor for 25 minutes at 2,25000 g. Late phagosomes were present at the interface between 25% sucrose and cytosol. Late phagosomes were collected with the cytosol, aliquoted (36 µl), and snap-frozen in liquid nitrogen.

2. Early Phagosomes

To isolate early phagosomes, a similar protocol was followed as the Late Phagosomes isolation with a few modifications. Synchronisation (4°C) was performed for 1 hr, and the pulse (22°C) time was reduced to 5 min without any chase. These cells were then collected, washed and lysed as written above. NOTE: The incubator temperature

(22°C) should be strictly maintained during cell growth in suspension and phagosome isolation to avoid cellular stress-induced effects on early phagosome maturation.

3. Post-nuclear Supernatant (PNS)

Cells grown in 100 ml suspension culture were pelleted down at 1000g ,4°C for 5 mins and washed in chilled Sorensen's buffer (pH=6.5). The cell pellet was weighed and lysed as described in the LP section above. The lysate was then centrifuged at a low speed of 2000g,4°C for 7 mins, and PNS was collected.

**Purification of Kinesin-1**

Fresh goat brain (15 g) was taken and lysed in 20 ml of homogenisation buffer (35 mM Pipes, 5 mM $MgSO_4$, 1mM EGTA, 0.5 mM EDTA, 1.5x PIC, 1mM DTT and 1 mM PMSF, pH adjusted using KOH to 7.2) and 0.5 mM ATP using Dounce. The homogenate was centrifuged for 1hr at 40,000 rpm,4°C and the high-speed supernatant (HSS) was collected. Then, 1 mM GTP and 20 µM taxol were added to the HSS, and native microtubules were polymerised at 37°C for 30 min in waterbath. Native microtubules were removed by pelleting in MLS 50 for 30 minutes at 150,000 g, 31°C. Then the supernatant was collected and incubated with microtubules (0.25mg/ml tubulin polymerised separately), hexokinase binding buffer (1 mM AMP-PNP, 10mg/ml glucose in homogenisation buffer), and 1 unit/ml hexokinase, with an additional 4 mM $MgSO_4$. This step was performed for 20 min at 30°C with 10-20 invert mixes to allow the motors to bind to the added microtubules. On the other hand, a two–step sucrose gradient (12.5% and 25%) was prepared with 1x PIC, 1 mM DTT, 0.2 mM GTP, and 20 µM Taxol in PMEE buffer (35 mM Pipes, 5 mM MgSO4, 1 mM EGTA, 0.5 mM EDTA). After incubation, we layered the binding mixture onto the sucrose cushions and centrifuged in MLS 50 for 50 minutes at 150,000 g, 31°C. Then the pellet was collected and dissolved in release buffer (1X PIC, 1mM DTT, 10mM ATP, 5mM $MgSO_4$ in PMEE) at 30°C in waterbath. Note that ATP was used to release the motor proteins from the microtubules. The release mixture was pelleted at 150,000g for 30 minutes, 31°C to remove the microtubules. Dynein-dynactin complex and kinesin-1 were separated from the release mixture based on the protocol described in [3,4]

**Isolation of Motor-rich fraction from *Dictyostelium***

0.36mg of tubulin was polymerised using a 1:1 (v/v) BGT (BRB 80 + 20 µM taxol + 10 mM GTP) at 37°C for 45 mins and kept separately. Dictyostelium cells were pelleted and washed twice using Sorensen's buffer. After washing, the pellet was lysed in LB-30 lysis buffer with protease inhibitors, as mentioned earlier. Note that the lysis was done in the absence of ATP. This lysate was subjected to a low-speed (1800g, 5 min) spin followed by a high-speed (2,25000 g, 25 min) spin on an LB25 sucrose gradient at 4°C. The high-speed supernatant (HSS) was collected after the spin. 15 U/ml Hexokinase, 4 mM glucose, polymerised microtubules, 20 µM taxol, 1.7mg/ml AMP-PNP, and 4 mM $MgCl_2$ were added to the HSS, which was then incubated on ice for 20 min. This step led to ATP depletion in the HSS and binding of motor proteins to the externally added microtubules. This mixture was centrifuged to pellet the microtubules at 1,20,000 g for 20 min at 4°C. The HSS after depletion of motor proteins was discarded. The pellet was collected and fully dissolved in the release buffer (1.5x PIC, 30 mM Tris pH 8, 4 mM EGTA, 15% sucrose, 10 mM ATP, 10 mM $MgCl_2$) for 30-45 mins on ice. This step allowed the motor proteins to detach from the microtubules upon ATP hydrolysis. Then the solution was centrifuged (1,80,000g) in TLA 100 for 20 mins at 4°C. The supernatant was collected, and the microtubule pellet (MT pellet) was discarded. The supernatant is the motor-rich fraction and can be used directly for motility.

For single-molecule experiments, Dictyostelium dynein-dynactin complex and DdUnc104 were separated from the motor-rich supernatant. Note that for separation, the release buffer was prepared containing 5% sucrose so that it could be loaded onto a 10–40% linear sucrose gradient in LB (30 mM Tris pH 8, 4 mM EGTA) and centrifuged at 1,42,000 g for 18 hrs, 4°C. Different fractions were collected and subjected to SDS-PAGE to assess the separation.

***In-Vitro* Motility Assay**

Coverslips were cleaned with acetone, dried under $N_2$, and subsequently treated with plasma to increase surface hydrophilicity. These hydrophilic coverslips were coated with Poly-L-lysine solution (500 µl PLL, 36ml ethanol, 4ml methanol) for 45 min and dried in a hot-air oven at 80°C. Flow chambers were made by sticking the PLL-coated coverslips on the glass slide at a 90° angle. The neat microtubules (as mentioned in the tubulin polymerisation step) or polarity labelled microtubules were diluted, flowed into the chamber, and left at room temperature for 20 mins. We visualised the microtubule on the coverslip surface of the flow chamber in DIC mode using a 100X oil-immersion objective on a custom-built optical trap [5]. Different cargoes were used (as described below) supplemented with

1mM ATP and the ATP regenerating system (20 mM MgCl2, 40 mM creatine phosphate, and 40 U/mL creatine kinase when needed) :

1. For bead motility assay, the beads were incubated with either purified kinesin-1, purified DdUnc104 or purified dynein for 5 minutes on ice. Motility buffer (30 mM Tris pH 8, 4 mM EGTA, 30% sucrose) and blocking buffer (1mg/ml casein and 10 µM Taxol) were added to the mixture for blocking of the coverslip surface. The final volume of the mixture was 36µl.
2. For SSLB motility assay, spherical-supported lipid bilayer cargoes [for preparation, refer to Kumar et al Cell Rep. 43 (2024)] were incubated with motor-rich fraction from Dictyostelium for 5 minutes on ice. Motility buffer and blocking buffer were added to the mixture. The final volume of the mixture was 36µl.
3. For endosome motility assay, the PNS collected was diluted (1:40) using LB-15 (30 mM Tris pH 8, 4 mM EGTA, 15% sucrose) before use.
4. For phagosome motility assay, one aliquot of 36µl isolated phagosomes was directly used.

The trap stiffness was using either $V_{match}$ or the power-spectra method [7]. Free or no-load motion of the different cargoes was observed using DIC microscopy. For in vitro stall experiments, we utilised the optical trap's ability to generate an opposing force on the cargoes towards its centre. This behaviour of the optical trap was modelled as a spring with a certain stiffness; hence, the opposing force is $F_{TRAP}$ = $K_{TRAP}$ * DISPLACEMENT. $K_{TRAP}$ was calculated for each cargo using the video-matching technique described in [7] to determine the stalling force. All the data were recorded using a customised LabView program and QPD setup described in our earlier papers [1,5,6].

**<u>Superstall Experiments</u>**

***IN-VITRO***

In Superstall experiments, we regulated the opposing force or 'load' on the cargo; by adjusting $K_{TRAP}$ or DISPLACEMENT. Catch bonds during dynein-MT interaction can be artificially increased by applying an opposing force higher than the stall force of the dynein ($F_{TRAP} > F_{DYNEIN}$) using the optical trap. For in vitro studies, we increased the DISPLACEMENT in the direction of the motion using a piezo-controlled stage to pull the cargo farther from the trap centre. This forces the dynein into a high-load regime. We termed these events Superstall, as shown in Figure 2D; the piezo's movement is indicated by a green arrow. A similar experiment was done by superstalling kinesin as a control.

***INSIDE CELLS***

For these studies, latex beads (750nm diameter) were phagocytosed into J774 cells maintained in a glass-bottom Petri plate. The spindle-shaped cells (see Supplementary Figure 4A) allowed us to identify the direction of motion: perinuclear is minus-directed, and peripheral is plus-directed. After 10 minutes of incubation, bidirectionally moving early phagosomes were observed and trapped inside the cell using a custom-developed LabView program. For the above described *in-vitro* studies, a freely floating EP is first trapped and brought down onto a polarity-labelled MT that is clearly visible and aligned horizontally. It is therefore easy to identify the direction (left or right) in which the piezo must be moved to cause a superstall for *in vitro* experiments. However, individual MTs are not visible, and the precise MT orientation is not known inside cells. Therefore, instead of moving the piezo to cause a superstall, we rapidly increased the laser power when an EP was moving against the trap inside cells. Based on our trap calibrations, this jump in laser power exerted an additional force of ~2 pN on the EP. Thus, in a typical superstall experiment inside cells the EP was allowed to stall at ~6pN force before an additional force of ~2 pN (superstall) was exerted by raising the laser power.

After a Stall or a Superstall force had been applied on an EP, we turned off the trap and observed the direction of motion of that EP. To stall/superstall dynein, we trapped the EP between minus-directed motions; whereas for kinesin, we trapped it between plus-directed motions. AVI videos of the entire sequence of events before and after stall/superstall was recorded. Note an important difference related to centering of the trap between the in-vitro and intracellular experiments. A trapped EP/LP may be attached to other invisible cellular structures, and may therefore never be really centered in the optical trap. One must test post-facto using the video tracks of EPs that the trapping position falls precisely at a point along the trajectory of a motile EP during force measurements. Typically, ~70% of the data is rejected because this condition does not hold. These issues have been discussed in detail earlier (Rai et al Cell 2013), where we have developed methods for quantitative force measurement on LPs inside J774 cells. Such precautions were employed when interrogating force measurement of bidirectional EPs inside cells. A bidirectionally moving EP was first identified inside a cell, the cursor was then brought over the position of the EP. A click of the mouse recorded the EP position, causing the piezo stage to displace the EP rapidly to the predetermined trap position, thus trapping the EP. The EP was now allowed to move against the trap to record stalls (no change in laser power), or superstalls (laser power increased; see earlier). A video recording of the entire event was made, using which the movement of the EP was later tracked at subpixel (~5 nm) resolution. The trajectory of these video tracks was analyzed to either reject (trap not centered; see earlier) or select an EP for further analysis. Fig 3C (see the yellow and blue boxes) shows representative X-Y trajectories of some EPs that were selected for analysis. Fig 3C (green box) also shows the video track of a purified EP moving *in-vitro* that stalled against the trap and then reversed direction along a single MT. It becomes clear by comparing the Y-displacements for data in green, yellow and blue boxes that EPs were also very likely reversing direction along the same MT inside cells.

**Monte Carlo Simulations**

Bidirectional cargo transport was simulated using the Gillespie Stochastic Simulation Algorithm (SSA) [8]. At each time step, one event (attachment, detachment, stepping or activation) was selected with probability proportional to its instantaneous rate. A fixed number of kinesin and dynein motors were coupled to a central cargo. Each motor was modelled as a linear elastic spring with stiffness $k_{motor}$ = 0.3 pN/nm and rest length $L_0$ (Supplementary Table 1), thereby implementing a condition of unequal load sharing by motors (10). The forward stepping rate was defined as the load-dependent velocity divided by the step size, backward stepping was prohibited. At every time step, the new equilibrium cargo position was determined by a simple force balance ($\sum F_i = 0,\ i \in Motor$) and the position of cargo and motor tails were subsequently updated

The force dependent velocity for both motors was given by [9,10]:

$$v(F) = \begin{cases} v_0 & if\ F < 0 \\ v_0 \left(1 - \frac{F}{F_{stall}}\right)^w & if\ 0 \leq F < F_{STALL} \\ 0 & if\ F\ \geq\ F_{STALL} \end{cases} \qquad (1)$$

Where $F$ is the instantaneous load on the motor (see Supplementary Table 1 for $F_{Stall}$). Here $v_0$ is the forward motor velocity (taken as 1 µm/sec for both motors). The exponent $w$ = 2.0 for kinesin and 0.5 for dynein, based on the force-velocity curves for these motors [5].

Here The MT-binding rate for each motor was taken as constant ($K_{ON}$ ; see Supplementary Table 1). The detachment rates followed different models depending on the simulation. The following table summarises the different models:

| Motor protein | Condition | Event | Modelled as |
|---|---|---|---|
| Kinesin | In all Simulations | Detachment | Slip Bond [9] |
| Dynein | No catch bond | Detachment | Slip Bond [9] |
| | Catch bond without Inactive state of dynein | Detachment | TFBD model [10] |
| | Catch bond with Inactive state of dynein | Detachment | TFBD model [10] |
| | | Entropic Constraint | See main text |

In the Slip Bond scenario, the detachment rate of the individual motor under load is given by [9,10]:

$$\varepsilon(F) = \epsilon_0 \cdot \exp\left[|F|/F_d\right] \qquad (2)$$

$F$ is the instantaneous load on each motor. $F_d$ is the detachment force scale (= 4 pN for kinesin and 0.67 pN for dynein [10]). $\epsilon_0$ is the no-load detachment rate (Supplementary Table 1).

In the Threshold Force Bond Deformation (TFBD) scenario, the dynein detachment rate for a single dynein exhibited two regimes depending on the threshold force (= 1.3 pN; see Ref. [10] of Supplementary Information). When the instantaneous load on the motor is below the trigger force, the detachment rate under load is given by equation 2 (slip-bind). Above the trigger force, the detachment rate under load is given by (see Ref. [10] of Supplementary Information):

$$\varepsilon(F) = \epsilon_0 \cdot \exp\left[-E_d(F) + (|F|/F_d)\right] \quad (3)$$

Deformation energy due to catch bond formation $-E_d(F) = \alpha \cdot \left(1 - \exp\left[-\frac{(|F|-1.3)}{f_0}\right]\right)$ (4)

Where $F$ is the instantaneous load on each dynein, $\alpha$ is the catch bond strength (= 58.08 $K_B T$), $f_0$ is the force scale (= 29 pN) that tunes the curvature of $\epsilon(F)$ in the catch bond region shown in Figure 1C. $F_d$ is the detachment force (= 0.67 pN for dynein), and $\epsilon_0$ is the no-load detachment rate.

Post-processing and segmentation of the simulated tracks

Simulated cargo trajectories were processed using a sliding-window velocity analysis. Raw positions, cumulative forces, and motor binding states were temporally interpolated to a step size of Δt = 0.01s. Mean velocity was computed from displacements over a 0.5s sliding window. Sub-trajectories were classified into three states using a velocity threshold $V_{thres}$ = 150 nm/sec. Segments with ($V > V_{thresh}$) were assigned to plus-directed runs, segments with ($V < -V_{thresh}$) were assigned to minus-directed runs, and intervening segments with $|V| \leq V_{thresh}$ were assigned to pauses or Tug-of-war [1]. After classification, contiguous segments shorter than 0.15 s [refer to 1] were merged into the preceding segment to suppress spurious events that would otherwise be indistinguishable in an experimental setting.

**Quantification and Statistical Analysis**

ImageJ was used to post-process the deep-SIM image of the J774 cell. Origin-2021 was used to prepare the graphs. Customised LabVIEW (National Instruments) programs were used to visualise and track optical trap-generated data. The fittings and Monte-carlo simulations were written in Python and uploaded to GitHub. Based on the normality of the dataset, either a two-tailed t-test or a two-tailed Mann-Whitney test was used to assess significance. The inferences were verified using a bootstrapping method for comparison of non-normal data sets (see Methods in main manuscript).

#### SUPPLEMENTARY VIDEO CAPTIONS

**Video S1.** Free motion (no optical trap) of bidirectionally moving EP inside a J774 cell, related to Figure 1A. The motion of the EP was tracked in the elongated region of the cell. The refractile latex bead EP (annotated by white arrow) moves in back -and- forth manner along a linear track.

**Video S2.** *In-vitro* motion of bidirectional EP purified from *Dictyostelium* against an optical trap, related to Figure 1F. Motion of EP is marked by the white arrow moving back-and-forth across the red line (trap centre). The microtubule is not in focus.

**Video S3.** Force-Induced reversal of motion from dynein-driven to kinesin-driven of an EP inside a J774 cell, related to Figure 3C. A motile dynein-driven EP (moving towards the nucleus, yellow arrow) was identified and was rapidly moved by a piezo stage to the known trap coordinates ($X_{TRAP}$, $Y_{TRAP}$) using a custom-developed LabVIEW Program. The motion of EP was restricted by the trap, as the SD in position was visibly reduced (denoted by the red arrow). The laser power was increased, as mentioned in the video, to superstall dyneins on the EP. The trap was turned off following which the EP showed a reversal in motion (kinesin-driven, i.e., away from the nucleus, denoted by the cyan arrow) along the same linear track as before.

**Video S4.** Force-Induced reversal of motion from kinesin to dynein on an EP inside J774 cell, related to Figure 3C. A motile kinesin-driven (i.e., away from the nucleus, denoted by the cyan arrow) EP was identified in real time and was rapidly displaced to the known trap coordinates. Further details as in Video S3.

### APPENDIX 1,2,3

Force Induces a Collective Inactivity in Dyneins to Auto-Regulate Bidirectional Transport of Vesicles. Dwiteeya Chaudhury et. al.

Note :- Citations in these Appendices pertain to the Reference List in Supplementary Information

#### Appendix 1: Mean Inactive Times

We consider the case of pure dynein driven motion. A cargo returns to the optical trap centre when all attached dyneins unbind from the microtubule (MT). This is the start of the inactive period. The inactive time $\tau$ is then defined as the time interval between the event of all dynein motors detaching from the MT and the event of the reattachment of the first unbound dynein to the MT.

In general, an unbound cargo with $N$ total dyneins within its contact area, $A_{CONTACT}$, will have a population of unbound dynein which are *inactive* ($D^*$) and rest which are *active* ($D$). The overall state of the unbound cargo, can be labeled in terms of total number of active motors, $m$ and total number of dyneins $N$, and is denoted as $(m, N)$, with $N - m$ then being the number of inactive ($D^*$) dyneins.

For such a state - $(m, N)$, two possible transitions can occur: any one of the motors in $D^*$ state converts to $D$. This process occurs with a rate $(N - m)k(m)$, where $k(m)$ is the activation rate. Alternately, any one of the motor in active state $D$, binds to the MT, with a rate $m\pi(m)$, thus ending the inactive period.

The activation rate $k(m)$ and the binding rate $\pi(m)$ can, in general, depend on the number of active motors. The limit $k(m) = k_o$ and $\pi(m) = \pi$ correspond to the no interaction case, where binding and activation events are independent of other motors. The mean inactive time, $T(m, N) = \langle\tau\rangle$ is then a mean first passage time from the state $(m, N)$ to the state with one bound dynein. The general expression of $T(m, N)$ can then be written as a set of recursion relations [11,12],

$$
\begin{aligned}
T(0,N) &= \frac{1}{Nk(0)} + T(1,N) \\
T(1,N) &= \frac{1}{(N-1)k(1)+\pi(1)} + \frac{(N-1)k(1)}{(N-1)k(1)+\pi(1)}\,T(2,N) \\
&\vdots \\
T(m,N) &= \frac{1}{(N-m)k(m)+m\pi(m)} + \frac{(N-m)k(m)}{(N-m)k(m)+m\pi(m)}\,T(m+1,N) \\
&\vdots \\
T(N-1,N) &= \frac{1}{k(N-1)+(N-1)\pi(N-1)} + \frac{k(N-1)}{k(N-1)+(N-1)\pi(N-1)}\,T(N,N) \\
T(N,N) &= \frac{1}{N\pi(N)}
\end{aligned}
$$

Let $R_m = (N-m)k(m) + m\pi(m)$. Then, solving the above set of equations self-consistently, we have,

$$T(m,N) = \sum_{i=m}^{N} \frac{1}{R_i} \prod_{j=m}^{i-1} \frac{(N-j)k(j)}{R_j}\,, \qquad 0 \le m \le N. \tag{1}$$

For the special case where the cargo unbinds with all motors inactive ($D^*$), the mean inactive time is given by,

$$T(0,N) = \sum_{i=0}^{N} \frac{1}{R_i} \prod_{j=0}^{i-1} \frac{(N-j)k(j)}{R_j} \tag{2}$$

##### 1.1 No interactions: Bounds of $T(0,N)$

While the above Eqs. 1, 2 provides the full analytical solution, it is also instructive to derive simple interpetative bounds for the no interaction case, where each dynein behaves independently of the others, $k(m) = k_o$ and $\pi(m) = \pi$. From the set of recursion relations, we have exactly,

$$T(0,N) = \frac{1}{Nk_0} + T(1,N),$$

where the first term corresponds to the mean time of the first dynein (out of $N$) becoming active ($D^* \to D$). Now, again from the recursion relations,

$$T(1,N) = \frac{1}{(N-1)k_o + \pi} + \frac{(N-1)k_o}{(N-1)k_o + \pi} T(2,N).$$

If binding is much faster than activation, $\pi \gg (N-1)k_0$, we can neglect the second term, and this reduces to $T(1,N) \approx 1/\pi$. Thus in this limit, we have,

$$T(0,N) \simeq \frac{1}{Nk_0} + \frac{1}{\pi}. \tag{3}$$

In the expression above, $T(0,N)$ is the mean inactive time when all the $N$ unattached dynein motors are inactive. The terms appearing in Eq. 3 maybe understood in the following manner. The first term corresponds to the mean time for the first dynein out of $N$ dyneins to become active, while the second term corresponds to the mean time for that active dynein to bind to the MT and thus ending the inactive period. Note that for any finite values of $k_o$, the true value of $T(1,N)$ is strictly smaller than $1/\pi$ value since there are additional transition of dynein from inactive to the active state. This in turns implies an increase in the effective binding rate, and consequent reduction of the inactive time. Thus Eq. 3 then gives an upper bound to the mean inactive time.

##### 1.2 Entropic Constraint Model

The Entropically Constrained Model (see Main text) that posits a steric hindrance which prevents the activation of an inactive dynein ($D^* \to D$) due entanglements. The per dynein activation rate is then modulated by the presence of other $D^*$ dyneins to yield an activation rate of the form,

$$k(m) = \frac{k_o}{1 + \Phi(N-m-1)}$$

where, $\Phi$ is an "entanglement parameter". A large $\Phi$ prevents the initiation of the activation cascade when starting from the all-inactive state, resulting in no active dynein available to bind to the MT. Thus hindered activation can lead to large increasing inactive times with increasing entanglement parameter $\Phi$.

### Appendix 2 Splitting Probabilities

Consider a cargo carried by $N$ dynein motors. At superstall, all $N$ motors are attached to the MT through an induced catch-bonded state, $D^{CB} + MT$ (see Fig. 2F in main manuscript). Once a catch-bonded motor unbinds, it is in the inactive conformation $D^*$, from which it may become active ($D$), and an active unbound motor may in turn rebind to the MT, re-entering the catch-bonded state $D^{CB} + MT$ directly, since a newly reattached motor shares the load with the others that are still attached. The cargo detaches from the MT at the instant when the last catch-bonded motor unbinds. At this instant, the $N$ unbound motors are distributed between the active and inactive conformations in some combination. The splitting probability $P(m, N)$ is defined as the probability that, starting from the state in which all $N$ motors are catch-bonded and bound to MT, the cargo detaches with exactly $m$ active (unbound) motors , while the remaining $N - m$ (unbound)motors are inactive [12,13]. Since the last motor to unbind is, by definition, still inactive at the instant of detachment (it has had no time to activate), $m$ ranges only over $0, 1, \ldots, N - 1$, and $\sum_{m=0}^{N-1} P(m, N) = 1$.

We denote the state of the system in terms of number of motors attached to MT ($L$), and the number of unattached motors which are *active* ($j$). Thus the state of the system maybe represented by a pair $(L, j)$. If $N$ is the total number of motors (attached and unattached) then $L \in \{0, 1, \ldots, N\}$, and $j \in \{0, 1, \ldots, N - L\}$. In general, for any particular state $(L, j)$, while $N - L$ unattached motors are active ($D$); the other $N - L - j$ unattached motors are inactive ($D^*$). The initial, fully catch-bonded state is $(N, 0)$. Once $L = 0$ the cargo has detached, and the states $j = 0, 1, \ldots, N - 1$ at $L = 0$ are the $N$ possible detachment conformations.

From a state $(L, j)$ with $L \geq 1$, three types of transition are possible:

- **Unbinding**: any one of the $L$ attached motors can unbind, at total rate $L\varepsilon$, taking $(L, j) \to (L - 1, j)$ (an unbound motor is always inactive immediately after unbinding);
- **Rebinding**: any one of the $j$ active unattached motors can rebind, at total rate $j\pi$, taking $(L, j) \to (L + 1, j - 1)$;
- **Activation**: any one of the $N - L - j$ inactive unattached motors can activate, at total rate $(N - L - j)k$, taking $(L, j) \to (L, j + 1)$.

Here $\varepsilon$, $\pi$, and $k$ are, respectively, the detachment rate of a catch-bonded motor, the attachment rate of an active unattached motor, and the activation rate of an inactive unattached motor. The total rate out of $(L, j)$ is therefore

$$R(L, j) = L\varepsilon + j\pi + (N - L - j)k.$$

Let $P(L, j; m)$ denote the probability that starting from an initial state $(L, j)$ where, $L$ motors are attached to MT and $j$ unattached active motors, the cargo eventually detaches from MT with exactly $m$ active motors. When $L = 0$, this corresponds to the situation where the initial state is the unbound state of cargo with no attached dynein. Therefore it follows that when $L = 0$,

$$P(0, j; m) = \begin{cases} 1, & j = m, \\ 0, & j \neq m. \end{cases}$$

For $L \geq 1$, conditioning on which of the three transitions out of $(L, j)$ occurs first (each is the first to occur with probability proportional to its rate, by the memoryless property of the competing exponential waiting times) gives the balance equation,

$$R(L, j)\, P(L, j; m) = L\varepsilon\, P(L-1, j; m) + j\pi\, P(L+1, j-1; m) + (N-L-j)k\, P(L, j+1; m). \quad (4)$$

The quantity of interest, the splitting probability itself, is $P(m,N) \equiv P(N,0;m)$. It maybe noted that $P(N,0:m)$ (and equivalently splitting probability $P(m,N)$ ) corresponds to the probability that the cargo unbinds from MT with $m$ unbound active motor, starting from an initial fully catchbonded configuration for which all $N$ motors are attached to MT (and none are unattached and active).

Formally, the transition graph on $(L,j)$ contains cycles rather than being a simple directed acyclic graph. For example, the set of transtions, $(L,j) \to (L,j+1) \to (L+1,j) \to (L,j+1)$, via activation, then rebinding, then unbinding, returns to a state already visited. Consequently the balance equations in Eq. 4 must be solved as a coupled linear system.

To formalize this, collect the $\sum_{L=1}^{N}(N-L+1) = N(N+1)/2$ transient states $(L,j)$, with $1 \le L \le N$ and $0 \le j \le N-L$, into a single index, and let $\mathbf{p}_m$ be the corresponding column vector of unknowns $P(L,j;m)$. The balance equations take the matrix form

$$\boxed{(D-A)\,\mathbf{p}_m \;=\; \mathbf{b}_m}$$

where $D = \mathrm{diag}\big(R(L,j)\big)$ is the diagonal matrix of total exit rates, $A$ is the matrix of transition rates between pairs of transient states (entries $L\varepsilon$, $j\pi$, or $(N-L-j)k$ for pairs connected by unbinding, rebinding, or activation respectively, and zero otherwise), and $\mathbf{b}_m$ collects the boundary contributions from transitions that land directly on $L=0$. The only such route is unbinding from a state $(1,j)$, which lands on $(0,j)$; the state $(1,j)$ therefore contributes an entry $\varepsilon\,\delta_{j,m}$ to $\mathbf{b}_m$, and this is the only way $\mathbf{b}_m$ can be nonzero.

Solving this linear system for each $m = 0,1,\ldots,N-1$ in turn, and reading off the entry corresponding to the initial state $(N,0)$, gives the full set of splitting probabilities $P(m,N)$, which by construction satisfy $\sum_{m=0}^{N-1} P(m,N) = 1$. For small $N$ this linear system is small enough to be solved explicitly by hand, as shown below.

##### 2.1 Two dyneins.

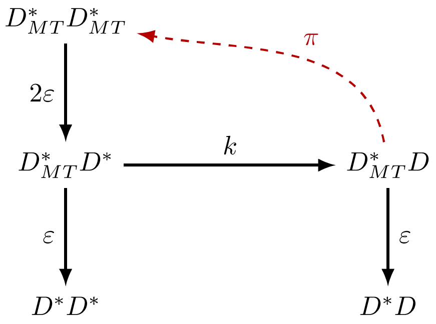


For $N=2$, the states are $(2,0) = D^*_{MT}D^*_{MT}$ (the initial state), $(1,0) = D^*_{MT}D^*$, $(1,1) = D^*_{MT}D$, and the two detachment outcomes $(0,0) = D^*D^*$ and $(0,1) = D^*D$.

The rates are $R(2,0) = 2\varepsilon$, $R(1,0) = \varepsilon + k$, $R(1,1) = \varepsilon + \pi$, and the balance equations read

$$\begin{aligned}
P(2,0;m) &= P(1,0;m),\\
P(1,0;m) &= \frac{\varepsilon}{\varepsilon+k}P(0,0;m) + \frac{k}{\varepsilon+k}P(1,1;m),\\
P(1,1;m) &= \frac{\varepsilon}{\varepsilon+\pi}P(0,1;m) + \frac{\pi}{\varepsilon+\pi}P(2,0;m).
\end{aligned}$$

Solving,

$$\text{prob.(unbinding through } D^*D^*) \equiv P(2,0;0) = \frac{\pi+\varepsilon}{k+\pi+\varepsilon},$$
$$\text{prob.(unbinding through } D^*D) \equiv P(2,0;1) = \frac{k}{k+\pi+\varepsilon}.$$

##### 2.2 Three dyneins.

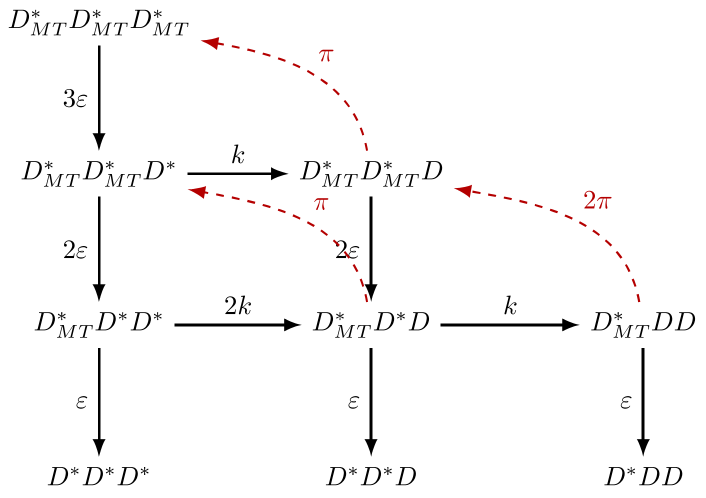


For $N = 3$, the states are $(3,0) = D^*_{MT}D^*_{MT}D^*_{MT}$ (the initial state), $(2,0) = D^*_{MT}D^*_{MT}D^*$, $(2,1) = D^*_{MT}D^*_{MT}D$, $(1,0) = D^*_{MT}D^*D^*$, $(1,1) = D^*_{MT}D^*D$, $(1,2) = D^*_{MT}DD$, and the three detachment outcomes $(0,0) = D^*D^*D^*$, $(0,1) = D^*D^*D$, $(0,2) = D^*DD$.

The rates are

$$R(3,0) = 3\varepsilon, \qquad R(2,0) = 2\varepsilon + k, \qquad R(2,1) = 2\varepsilon + \pi,$$
$$R(1,0) = \varepsilon + 2k, \qquad R(1,1) = \varepsilon + \pi + k, \qquad R(1,2) = \varepsilon + 2\pi,$$

and the balance equations read

$$P(3,0;m) = P(2,0;m),$$
$$P(2,0;m) = \frac{k}{2\varepsilon+k}P(2,1;m) + \frac{2\varepsilon}{2\varepsilon+k}P(1,0;m),$$
$$P(2,1;m) = \frac{2\varepsilon}{2\varepsilon+\pi}P(1,1;m) + \frac{\pi}{2\varepsilon+\pi}P(3,0;m),$$
$$P(1,0;m) = \frac{2k}{\varepsilon+2k}P(1,1;m) + \frac{\varepsilon}{\varepsilon+2k}P(0,0;m),$$
$$P(1,1;m) = \frac{k}{\varepsilon+\pi+k}P(1,2;m) + \frac{\varepsilon}{\varepsilon+\pi+k}P(0,1;m) + \frac{\pi}{\varepsilon+\pi+k}P(2,0;m),$$
$$P(1,2;m) = \frac{2\pi}{\varepsilon+2\pi}P(2,1;m) + \frac{\varepsilon}{\varepsilon+2\pi}P(0,2;m).$$

Solving, we obtain,

$$\text{prob.(unbinding through } D^*D^*D^*) = \frac{2\varepsilon^3 + 2\varepsilon^2 k + 7\varepsilon^2\pi + \varepsilon k\pi + 7\varepsilon\pi^2 + 2k\pi^2 + 2\pi^3}{\mathcal{N}}$$

$$\text{prob.(unbinding through } D^*D^*D) = \frac{k(\varepsilon + 2\pi)(5\varepsilon + 2k + 2\pi)}{\mathcal{N}}$$

$$\text{prob.(unbinding through } D^*DD) = \frac{k^2(5\varepsilon + 2k + 2\pi)}{\mathcal{N}}$$

with

$$\mathcal{N} \equiv 2\varepsilon^3 + 7\varepsilon^2 k + 7\varepsilon^2\pi + 7\varepsilon k^2 + 13\varepsilon k\pi + 7\varepsilon\pi^2 + 2k^3 + 6k^2\pi + 6k\pi^2 + 2\pi^3 :$$

##### 2.3 General trend with $N$

The general structure above lets $P(m, N)$ be computed for any $N$ by solving the corresponding linear system numerically, even for state-dependent activation rates. We use the standard no-load attachment rate, $\pi = 2.5/s$ and the experimentally measured single dynein activation rate $k_0 = 2.5/s$. The detachment rate of a catch-bonded dynein in general, depends on the force, and has a catch-ideal behaviour in the post-stall regime. This varies over a regime of $4.5/s$ immediately post-stall, to $\sim 1/s$ at large forces (see Fig. 1c). We assume an average detachment rate of $\varepsilon = 2/s$ in this catch-bonded state to reflect both the larger forces induced by the superstall experiments, and subsequent detachment of catchbonded dyneins, which increases the per-dynein force for the remaining catchbonded dyneins. Table 1 lists the probability $P(0, N)$ of detaching with *all* $N$ dyneins inactive, for $N = 1$ through 6, under both the no interaction model, and the entropically constrained model. As can be seen from the table, for non-interacting dyneins, the probability of detaching with all dyneins inactive rapidly falls with increasing dynein number. Conversely, collective inactivation through entropic constraints leads to a higher probability of detaching in the all-inactive state, with at 75% probability of detaching with all motors inactive for $N = 5$.

| $N$ | 1 | 2 | 3 | 4 | 5 | 6 |
|---|---|---|---|---|---|---|
| $P(0, N)$, no interaction | 1.000 | 0.643 | 0.361 | 0.189 | 0.095 | 0.046 |
| $P(0, N)$, entropic constraint | 1.000 | 0.643 | 0.683 | 0.729 | 0.754 | 0.768 |

Table 1: Probability $P(0, N)$ that the cargo detaches with all $N$ dyneins inactive, for $N = 1, \ldots, 6$, under no interaction ($\Phi = 0$) and entropic constraint ($\Phi = 4.54$), with $\varepsilon = 2.0/s$, $\pi = 2.5/s$, $k_0 = 2.5/s$ in both cases.

### Appendix 3 Distribution of Inactive Times

In Appendix 1, we had derived the expression for mean Inactive time of the cargo before it reattaches to MT. The inactive period terminates with the attachment of any of the active dynein to the MT. These states, where any one dynein attaches to MT ($D^{CB} + MT$), are the *absorbing states* for this process. We now proceed to determine the full distribution of the inactive time.

We define $S_m(t)$, the survival probability, as the probability that the cargo has not reached any absorbing state by time $t$, having started with $m$ active dyneins at $t = 0$. As before, a cargo with $m$ active dyneins can either undergo transition to a state with $m+1$ active dyneins with a rate $(N-m)k(m)$, or bind to the MT with a rate $m\pi$. Since the inactive period ends with the binding of a dynein to a MT, this second transition is a transition to an absorbing state. As soon as any of the active dyneins bind to the MT, cargo motion restarts, ending the inactive period. The backward master equation for $S_m(t)$ is [11]

$$\frac{dS_m(t)}{dt} = \sum_{m'} \omega_{m\to m'} S_{m'} - \omega_{m\to m'} S_m = -\big[(N-m)k(m) + m\pi\big] S_m + (N-m)k(m)\, S_{m+1}$$

(since any absorbing state has $S_{\text{abs}} = 0$ by definition). Writing $R_m \equiv (N-m)k(m) + m\pi$, as before, we have

$$\frac{dS_m}{dt} = -R_m S_m + (N-m)k(m)\, S_{m+1}(t), \qquad S_m(t=0) = 1.$$

Define the Laplace transform, $\tilde{S}_m(s) = \int_0^\infty e^{-st} S_m(t)\, dt$. The backward equation becomes

$$s\tilde{S}_m(s) - 1 = -R_m \tilde{S}_m(s) + (N-m)k(m)\, \tilde{S}_{m+1}(s),$$

giving the recursion

$$\tilde{S}_m(s) = \frac{1 + (N-m)k(m)\, \tilde{S}_{m+1}(s)}{s + R_m}.$$

At $m = N$, $R_N = N\pi$ and there is no activation term, so

$$\tilde{S}_N(s) = \frac{1}{s + R_N} = \frac{1}{s + N\pi} \qquad \Longrightarrow \qquad S_N(t) = e^{-N\pi t}.$$

We want to invert the recursion for a general $m$. Note that starting from the state $m$, the system passes through a strictly increasing sequence of states $m, m+1, \ldots$ before eventually being absorbed by a binding event at some level. At each level $l$ it visits, the memoryless holding time before the next event (either advancing to $l+1$ or being absorbed) is exponentially distributed with rate $R_l = (N-l)k(m) + l\pi$, the total rate of leaving state $l$. A holding time of rate $R_l$ contributes, in Laplace space, a single simple pole at $s = -R_l$. Thus $\tilde{S}_m(s)$ must be a sum of simple poles at $s = -R_m, -R_{m+1}, \ldots, -R_N$,

$$\tilde{S}_m(s) = \sum_{j=m}^{N} \frac{A_j^{(m)}}{s + R_j},$$

where the residues $A_j^{(m)}$ are unknown.

Since $\mathcal{L}^{-1}\{1/(s + R_j)\} = e^{-R_j t}$, inverting the transform therefore always produces $S_m(t)$ as a superposition of decaying exponentials $e^{-R_j t}$. Thus the total exit rates $R_j$ set the characteristic timescales $\tau_j = 1/R_j$ of the survival probability. This is true for both entropically constrained activation and the non-interacting case, with the rates $R_j$ suitably redefined to account for the specific dependence of the binding/activation rates on the number of active motors. Thus we have,

$$S_m(t) = \sum_{j=m}^{N} A_j^{(m)}\, e^{-R_j t}. \tag{5}$$

Unrolling the recursion for $\tilde{S}_m(s)$ fully in Laplace space, and then matching residues via a full multi-pole partial-fraction decomposition of the result, we obtain,

$$A_j^{(m)} = \left\{ \sum_{r=j}^{N} \frac{\prod_{l=m}^{r-1} (N-l)k(l)}{\prod_{l=m,\, l\neq j}^{r} (R_l - R_j)} \right\} \tag{6}$$

Together, Eqs. 5 and 6 define the survival probability distributions.

The cumulative distribution function for first passage times, $F_m(t)$ denotes the probability that inactive period has terminated at any time $\tau < t$, and hence is simply given by $F_m(t) = 1 - S_m(t)$. The probability distribution of first-passage times (inactive times) is then

$$p_m(\tau) = -\left.\frac{\partial S_m(t)}{\partial t}\right|_{t=\tau}.$$

The mean inactive time is

$$\langle \tau_m \rangle = \int_0^\infty d\tau\, \tau\, p_m(\tau) = \int_0^\infty d\tau\, S_m(\tau) = \tilde{S}_m(s=0),$$

which recovers exactly the hierarchy of MFPT equations, $T(m, N)$, derived in Appendix 1.

##### 3.1 No interactions

In this case, with $k_0 = \pi$, all $R_l = Nk_0 = \rho$ (say), and so this is the totally confluent case with all poles conincident. In this case, we have,

$$\tilde{S}_m(s) = \sum_{r=m}^{N} \left[ \prod_{l=m}^{r-1} (N-l)k_0 \right] \frac{1}{(s+\rho)^{r-m+1}} = (N-m)! \sum_{n=1}^{N-m+1} \frac{k_0^{n-1}}{(N-m+1-n)!} \cdot \frac{1}{(s+\rho)^n},$$

Inverting term by term, we have,

$$S_m(t) = (1 + k_0 t)^{N-m} e^{-Nk_0 t}. \tag{7}$$

This is then the survival probability for the non-interacting case.

##### 3.2 Entropic Constraint Model

For entropically constrained activation, with $k_0 = \pi$, $(N - m - 1) = 0$ at $m = N - 1$, so $k(N-1) = k_0 = \pi$ identically, and therefore $R_{N-1} = R_N$. We thus have a single double pole at $R_{N-1} = R_N \equiv$

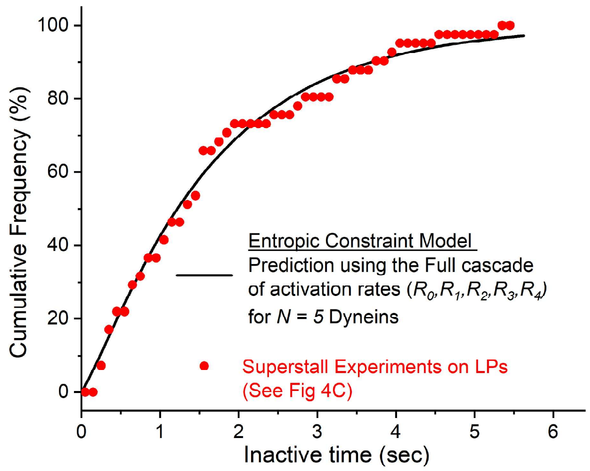


Appendix 3, Figure 1: Comparison of the distribution of inactive times for $N = 5$ dyneins from the theory, Eq. 8, with the superstall experimental data on LPs. Note that the theory has no free parameters, once $\Phi$ was determined from the mean inactive times.

$\rho$, with all other poles simple. In this case, one can show,

$$B_2^{(m)} = \left[\prod_{l=m}^{N-1}(N-l)k(l)\right]\prod_{l=m}^{N-2}\frac{1}{R_l-\rho},$$

$$B_1^{(m)} = \left[\prod_{l=m}^{N-2}(N-l)k(l)\right]\left[\prod_{l=m}^{N-2}\frac{1}{R_l-\rho}\right]\left[1-k_0\sum_{l=m}^{N-2}\frac{1}{R_l-\rho}\right],$$

and the survival probability is given by,

$$S_m(t) = \sum_{j=m}^{N-2} A_j^{(m)}\, e^{-R_j t} + \left[B_2^{(m)}\, t + B_1^{(m)}\right] e^{-\rho t}, \tag{8}$$

with $A_j^{(m)}$ as in Eq. 6. This full distribution matches remarkably well to the experimental data, as shown in Appendix-3, Fig. 1.

#### REFERENCES IN SUPPLEMENTARY INFORMATION

**Includes citations referred to in Appendix 1,2,3**


1. Soppina V, Rai AK, Ramaiya AJ, Barak P, Mallik R (2009) Tug-of-war between dissimilar teams of microtubule motors regulates transport and fission of endosomes. Proc Natl Acad Sci U S A 106(46):19381–19386.
2. Soppina, V., Rai, A., & Mallik, R. (2009). Simple non-fluorescent polarity labeling of microtubules for molecular motor assays. *BioTechniques*, *46*(7), 543–549.
3. Bingham, J. B., King, S. J., & Schroer, T. A. (1998). Purification of dynactin and dynein from brain tissue. *Methods in enzymology*, *298*, 171–184.
4. Schroer, T. A., & Sheetz, M. P. (1991). Two activators of microtubule-based vesicle transport. *The Journal of cell biology*, *115*(5), 1309–1318.
5. Rai, A. K., Rai, A., Ramaiya, A. J., Jha, R., & Mallik, R. (2013). Molecular adaptations allow dynein to generate large collective forces inside cells. *Cell*, *152*(1-2), 172–182.
6. Kumar P, Chaudhury D, Sanghavi P, Meghna A, Mallik R (2024) Phosphatidic acid-dependent recruitment of microtubule motors to spherical supported lipid bilayers for in vitro motility assays. Cell Rep 43(6).
7. Barak P, Rai A, Rai P, Mallik R (2013) Quantitative optical trapping on single organelles in cell extract. Nat Methods 10(1):68–70.
8. Gillespie, D. T. (1977). Exact stochastic simulation of coupled chemical reactions. *The journal of physical chemistry*, *81*(25), 2340-2361.
9. Müller, M. J., Klumpp, S., & Lipowsky, R. (2008). Tug-of-war as a cooperative mechanism for bidirectional cargo transport by molecular motors. *Proceedings of the National Academy of Sciences of the United States of America*, *105*(12), 4609–4614.
10. Puri P, et al. (2019) Dynein catch bond as a mediator of codependent bidirectional cellular transport. Phys Rev Res., *1*(2), 023019.
11. N.G. van Kampen. Stochastic Processes in Physics and Chemistry. Elsevier Science Publishers, Amsterdam, 1992.
12. Srividya Iyer-Biswas and Anton Zilman. First-Passage Processes in Cellular Biology. John Wiley Sons, Ltd, 2016.
13. Sidney Redner. A Guide to First-Passage Processes. Cambridge University Press, Cambridge, UK, 2001.
